# Quorum sensing inhibits Type III-A CRISPR-Cas system activity through repressing positive regulators SarA and ArcR in *Staphylococcus aureus*

**DOI:** 10.1101/2023.01.17.524377

**Authors:** Yang Li, Yuanyue Tang, Xiaofei Li, Nina Molin Høyland-Kroghsbo, Hanne Ingmer, Xinan Jiao, Qiuchun Li

**Author notes:** Correspondence (H. I.), (X. J.), (Q. L.). These authors contributed equally (Yang Li, Yuanyue Tang, Xiaofei Li).

## Abstract

CRISPR-Cas is an adaptive immune system that protects prokaryotes from the invasion of foreign genetic elements. The components and immunity mechanisms of CRISPR-Cas have been extensively studied, but the regulation of this system in *Staphylococci* remains unclear. Here, we show that in the *S. aureus* Type III-A CRISPR-Cas system, the P*cas* of 300 bp located in *cas1* displays as a critical regulatory node that initiates the transcription of *cas* gene clusters. We discovered two transcriptional regulators, SarA and CRP-like ArcR, promote the expression and activity of the CRISPR-Cas system by directly binding to the novel promoter P*cas*. Furthermore, we demonstrated that the cell-cell communication, known as quorum sensing (QS), inhibits the activity of CRISPR-Cas system by repressing the positive regulators SarA and ArcR. Bioinformatic analyses suggest that P*cas* is conserved in many Type II and III CRISPR-Cas systems in Firmicutes. Our data reveal a new regulatory mechanism for QS-mediated repression of the Type III-A CRISPR-Cas system, which may allow *S. aureus* to acquire foreign genetic elements encoding antibiotic resistance or virulence factors specifically at high cell density.

## INTRODUCTION

Bacteria have evolved multiple lines of innate and adaptive defenses to reduce the risk imposed by foreign genetic elements.^1^ The CRISPR-Cas system is an adaptive immune system in archaea and bacteria that defends against the invasion of foreign nucleic acids, such as viruses and plasmids.^2,3^ CRISPR-Cas systems have been classified into six types (I-VI) with 33 subtypes.^4^ Types I, III, and IV are composed of multi-subunit effector complexes, whereas Types II, V, and VI harbor a single effector protein that cleaves target nucleic acids.^4^ The defense processes of CRISPR-Cas systems consist of three distinct stages: adaptation, CRISPR RNA (crRNA) biogenesis, and interference.^5^ During adaptation, the invading nucleic acids derived from plasmids or bacteriophage (phage) viruses are recognized and incorporated into CRISPR arrays.^6,7^ The CRISPR array is transcribed into a long precursor CRISPR RNA (pre-crRNA), which is subsequently processed into mature crRNAs.^8,9^ Finally, the Cas proteins in combination with mature crRNAs form an effector complex to degrade the invading genetic elements.^10^ Unlike other CRISPR-Cas types, the Type III CRISPR-Cas system requires transcription of target sequences recognized by crRNAs for effective immunity.^11-13^ The target RNA binding to the effector complex triggers activation of the Cas10 HD and palm domains. The activated HD domain can non-specifically cleave ssDNA,^14-16^ and the palm domain is responsible for synthesis of cyclic oligoadenylates, which act as a secondary messenger to activate Csm6 or Csx1 RNase for non-specific RNA degradation.^17,18^

The Type III-A CRISPR-Cas system has been discovered at a frequency of 6.8% in *Staphylococci* species, including *S. aureus* and *S. epidermidis*.^3,19-21^ Furthermore, this system has been demonstrated to be functional in protecting against plasmid invasion and phage infection in both *S. aureus* and *S. epidermidis*.^3,12,20^ Although the functions and immune mechanisms of the Type III-A CRISPR-Cas system have been well studied, the transcriptional regulation of this system is not well understood.

Previous studies have demonstrated that CRISPR-Cas is positively regulated by QS in some Gram- negative bacteria.^22,23^ QS is a cell-cell communication process that allows bacteria to regulate their gene expression in response to cell density by releasing and detecting chemical signal molecules known as auto-inducers. At high cell density, when the auto-inducer concentration reaches a critical threshold, QS receptors are activated and turn on genes involved in group behaviors, such as virulence factors and biofilm formation.^24^ In Gram-negative bacteria, N-acyl-L-homoserine lactone (AHL) autoinducers are generally produced by LuxI type synthases and sensed by LuxR family transcriptional regulators. LuxI/R-type QS systems activate *cas* gene expression in the Gram-negative bacterium *Burkholderia glumae*.^25^ Additionally, CRISPR-Cas interference and CRISPR adaptation is activated in response to LuxR/I QS in both *Serratia* sp. ATCC39006 and *Pseudomonas aeruginosa*.^22,23^ At high cell density, phages can spread rapidly in the bacterial population.^26^ Presumably, QS activation of CRISPR-Cas protects bacteria specifically at high cell density. In Gram-positive *S. aureus*, QS is mediated via autoinducing peptides that are expressed from the *agr* locus and sensed by a two-component signal transduction system composed of a sensory histidine kinase, AgrC and a transcriptional response regulator, AgrA.^27^ The *agr* operon consists of the *agr*BDCA signaling cassette and RNA III gene, which are under the control of the P2 and P3 promoters, respectively.^28^ At high cell density AgrA induces the expression of RNA III, which in turn controls the expression of many key virulence factors, such as α-hemolysin.^29^ However, it is currently unknown whether QS regulates CRISPR-Cas systems in Gram-positive bacteria.

In this study, we identified a promoter, P*cas*, located in the coding region of *cas1* that initiates the transcription of *cas* gene clusters of the Type III-A CRISPR-Cas system. To dissect molecular mechanisms of *cas* genes regulation, we identified two regulators, SarA and cAMP receptor protein (CRP)-like ArcR, promote expression of *cas* genes by directly binding to P*cas*. Furthermore, we demonstrated that the QS response regulator AgrA indirectly represses *cas* genes expression by negatively regulating *sarA* and *arcR*. Thus, at high cell density, AgrA-mediated repression of *sarA* and *arcR* results in decreased *cas* genes expression and CRISPR-Cas-mediated interference activity in *S. aureus*.

## RESULTS

### P*leader* sequence drives the transcription of CRISPR1 array

To identify regulators of the Type III-A CRISPR-Cas system in *S. aureus*, we used the clinical methicillin-resistant *Staphylococcus aureus* (MRSA) strain TZ0912 carrying a complete Type III-A CRISPR-Cas system with two CRISPR arrays as the model (Figure 1A).^20^ We initially sought to discover direct CRISPR-Cas regulators through DNA pull-down using the promoter of *cas* gene clusters as bait. However, the promoter of the Type III-A CRISPR-Cas system has not been identified in *S. aureus*. Accordingly, we first determined whether the promoter for the *cas* genes transcript was located in the leader sequence upstream of CRISPR1. We chose to focus on CRISPR1 because this array locates upstream of *cas* gene clusters. We performed RT-PCR using nine sets of primers to amplify regions spanning different gene combinations (Table S5) to examine the potential co- transcription of the CRISPR1 locus and *cas* genes (Figure 1A). Each targeted region showed a specific size of DNA fragment from both gDNA and cDNA templates, indicating that the CRISPR1 locus and *cas* genes were transcribed on a single polycistronic mRNA by a potential promoter located in the leader sequence (Figure S1A). The leader sequence of the CRISPR1 locus was cloned into a promoterless *lacZ* fusion vector to evaluate its importance for transcription. We observed a 10-fold increase in the expression of the P*leader*-*lacZ* transcriptional fusion compared to the promoterless *lacZ* fusion, revealing that the leader sequence might harbor a key element driving transcription (Figure 1B). After construction of a Δ*leader* mutant, we evaluated the importance of the leader sequence for transcription of the CRISPR-*cas* array by qRT-PCR of *cas* gene expression. The expression of CRISPR1 pre-crRNA was downregulated 13-fold in the Δ*leader* mutant compared to that in the WT strain (Figure 1C), indicating that the leader sequence contained a promoter for CRISPR-*cas* in the WT. However, compared to the dramatically decreased level of *crispr1*, the abundance of *cas10* and *csm3* transcripts was only modestly downregulated (approximately 1.5-fold) in the Δ*leader* mutant (Figure 1C), implying that there was potentially a second promoter regulating *cas* genes expression. These results suggest that the leader sequence could drive the transcription of *crispr1* but not *cas* genes.

**Figure 1.**
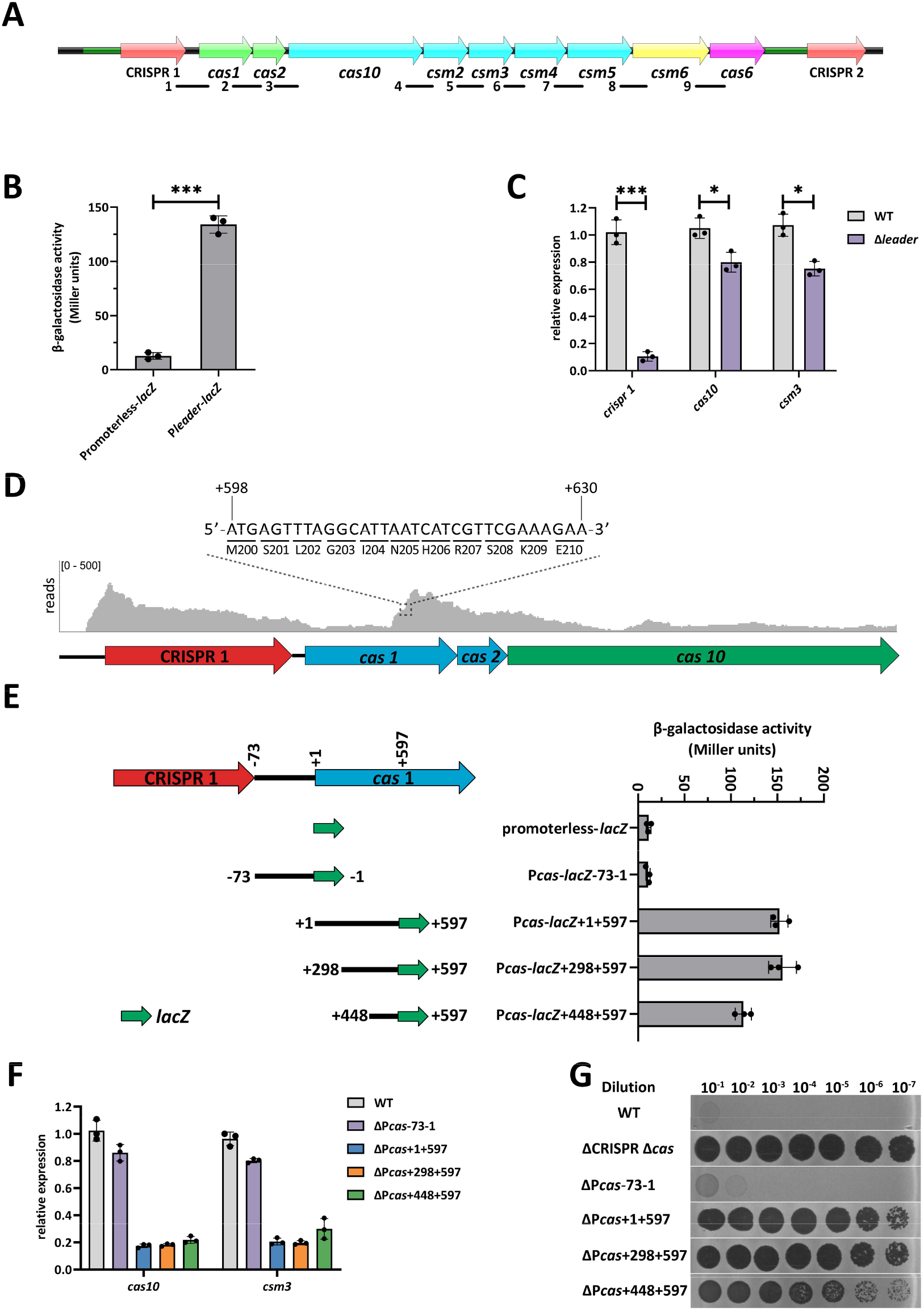
The leader sequence and a promoter located within the *cas1* gene promote transcription of CRISPR1 and *cas*, respectively. (A) Schematic of Type III-A CRISPR-Cas system in *S. aureus* TZ0912. The nine fragments (number 1 to 9) corresponding to intergenic regions and amplified in the RT-PCR experiments are indicated below. The leader of CRISPR 1 and CRISPR 2 array were indicated as dark green lines. (B) β-galactosidase activity of a promoterless *lacZ* fusion and of the P*leader*-*lacZ* reporter fusion in the WT. (C) qRT-PCR of *crispr, cas10* and *csm3* expression in the WT and in a Δ*leader* mutant, respectively. (D) RNA-seq reads were mapped to the WT genome to determine the relative abundance of crRNA from the CRISPR 1 array, *cas1, cas2*, and *cas10* by IGV software. An ATG at the (position between +598 to +600) is located within the *cas1* encoding region. (E) Transcriptional fusions are represented as lines with a short green arrow at their 3’ ends, indicating the respective genomic region (lines) and the *lacZ* reporter gene (green arrows). All of the positions indicated are relative to the translational start site (+1) of *cas1*. β-galactosidase activity of a promoterless *lacZ* reporter and of the respective *lacZ* reporter fusions were measured in the WT. (F) qRT-PCR of *cas10* and *csm3* in the WT and respective P*cas* mutants. (G) Tenfold serial dilution of phage phiIPLA-RODI was spotted on the bacterial lawns of *S. aureus* WT and mutant strains carrying the pCRISPR plasmid that expresses an additional crRNA for enhanced immunity. Plaquing experiments were replicated three times and consistent results were seen. All the data (B-C, E-F) are means ± standard deviation of three independent experiments. A two-tailed unpaired Student’s *t*-test was used to calculate *P* values; \**p*<0.05, \*\*\**p*<0.001.

### P*cas* sequence serves as a promoter of *cas* gene clusters

Next, we quantified the abundance of *cas* transcripts in the TZ0912 strain using RNA-seq to determine the predominant *cas* gene promoter. Within the *cas1* coding sequence, increased transcriptional activity was observed starting from the position of *cas1* +598 (Figure 1D). Further analysis showed that the position from +598 to +600 of *cas1* is ATG, indicating that there is a potential promoter upstream of position +597 in *cas1* to initiate the transcription of other *cas* genes (Figure 1D). To identify core promoters within this region, we constructed a series of transcriptional fusions carrying different 5’ end deletions and tested the constructs using the β-galactosidase assay (Figure 1E). Since there is a 73 bp fragment between the CRISPR1 array and *cas1*, we also fused it into the *lacZ* reporter gene (Figure 1E). The promoterless *lacZ* and P*cas*-*lacZ*-73-1 yielded minimal reporter activity. P*cas*-*lacZ*+1+597 resulted in a 15-fold increase in reporter activity compared to the promoterless *lacZ*. A further deletion of the remaining 300 bp at the 3’ end (P*cas*-*lacZ*+298+597) yielded a similar reporter expression level to that of P*cas*-*lacZ*+1+597, while P*cas*-*lacZ*+448+597 showed a slight decrease in reporter activity relative to P*cas*-*lacZ*+1+597 (Figure 1E). This suggests that a potential core promoter exists between positions +298 and +597 of *cas1*. To examine whether the putative promoters have an effect on *cas* genes transcription, we deleted the corresponding regions (Figure S1B) and quantified *cas* genes transcripts using qRT-PCR. Both *cas10* and *csm3* expression levels were reduced 5-fold in the ΔP*cas*+1+597, ΔP*cas*+298+597, and ΔP*cas*+448+597 mutants compared with those in the WT strain, whereas their expression was not significantly affected in the ΔP*cas*-73-1 mutant (Figure 1F). A strain lacking the 308 bp at the 3’-end of *cas1* (Δ*cas1*+598+905) had no effect on *cas* gene transcription (Figure S1B and S1C), further confirming that an active promoter is located between +1 and +597 of *cas1*.

To test the effect of putative promoters on CRISPR-Cas activity, we carried out plaque assays. Because our previous study showed that the WT strain only has weak immunity against phage philPLA-RODI infection in a plaque assay relative to the ΔCRISPR Δ*cas* mutant,^20^ we constructed a pCRISPR plasmid expressing a crRNA targeting phage philPLA-RODI, in order to enhance the immunity of the WT strain. Specifically, we designed a plasmid to carry the leader sequence of the CRISPR array, two repeats interrupted with a spacer targeting the phage philPLA-RODI *gp036* gene (Figure S1D); and the recombinant plasmid was transformed into the WT, ΔCRISPR Δ*cas*, and four promoter mutant strains (ΔP*cas*-73-1, ΔP*cas*+1+597, ΔP*cas*+298+597, and ΔP*cas*+448+597). Transformants were used as recipients when infected with phage philPLA-RODI. In accordance with our *cas* genes expression data, the EOP of phage philPLA-RODI on ΔP*cas*+1+597 and ΔP*cas*+298+597 mutant strains decreased by more than 10^6^-fold compared with WT (Figure 1G), whereas no difference in EOP was observed for ΔP*cas*-73-1 and Δ*cas1*+598+905 mutant strains compared with the WT (Figure 1G and S1E). Notably, the EOP on the ΔP*cas*+448+597 mutant was 10^3^-fold higher than that on the ΔP*cas*+298+597 mutant, which was consistent with the weak reporter activity detected in the β-galactosidase assay. These data demonstrate that an active promoter located between positions +298 and +597 of *cas1* (termed P*cas*) drives transcription of *cas* genes downstream of *cas1*.

### SarA and ArcR directly activate Type III-A CRISPR-Cas expression

To identify the direct transcriptional regulators of the Type III-A CRISPR-Cas system, we applied a DNA pull-down assay with the P*leader* and P*cas* sequences as bait to capture proteins from WT cellular lysates. Mass spectrometry analysis showed that 17 and 50 proteins had potential interactions with the P*leader* and P*cas* sequences, respectively (Table S1 and S2). Among the 17 proteins interacting with the P*leader* sequence, six proteins were predicted to be transcriptional regulators and 20 out of 50 proteins were potential transcriptional regulators that bind to the P*cas* sequence. In addition, four transcriptional regulators, SarA, SarR, ArcR, and Rot, were captured by both the P*leader* and P*cas* sequences in the DNA pull-down assays (Table S1 and S2).

To examine whether these four transcriptional regulators are involved in the transcription of both spacers and *cas* genes, we measured the expression levels of *crispr1, crispr2, cas10*, and *csm3* in four individual mutants (Δ*sarA*, Δ*sarR*, Δ*arcR*, and Δ*rot*) using qRT-PCR. In comparison with the WT, a marked increase in the expression of *crispr1* was observed in the Δ*sarA* mutant, whereas the expression of *cas* genes was decreased by 2-fold (Figure 2A). We observed increased expression of *crispr1* and *cas* genes in the Δ*sarR* mutant relative to the WT, but decreased expression in the Δ*arcR* mutant (Figure 2A). Furthermore, deletion of *rot* had no significant effect on the expression of *crispr1* and *cas* genes (Figure 2A). The expression of *crispr2* was not altered in any of the four mutants (Figure S2A).

**Figure 2.**
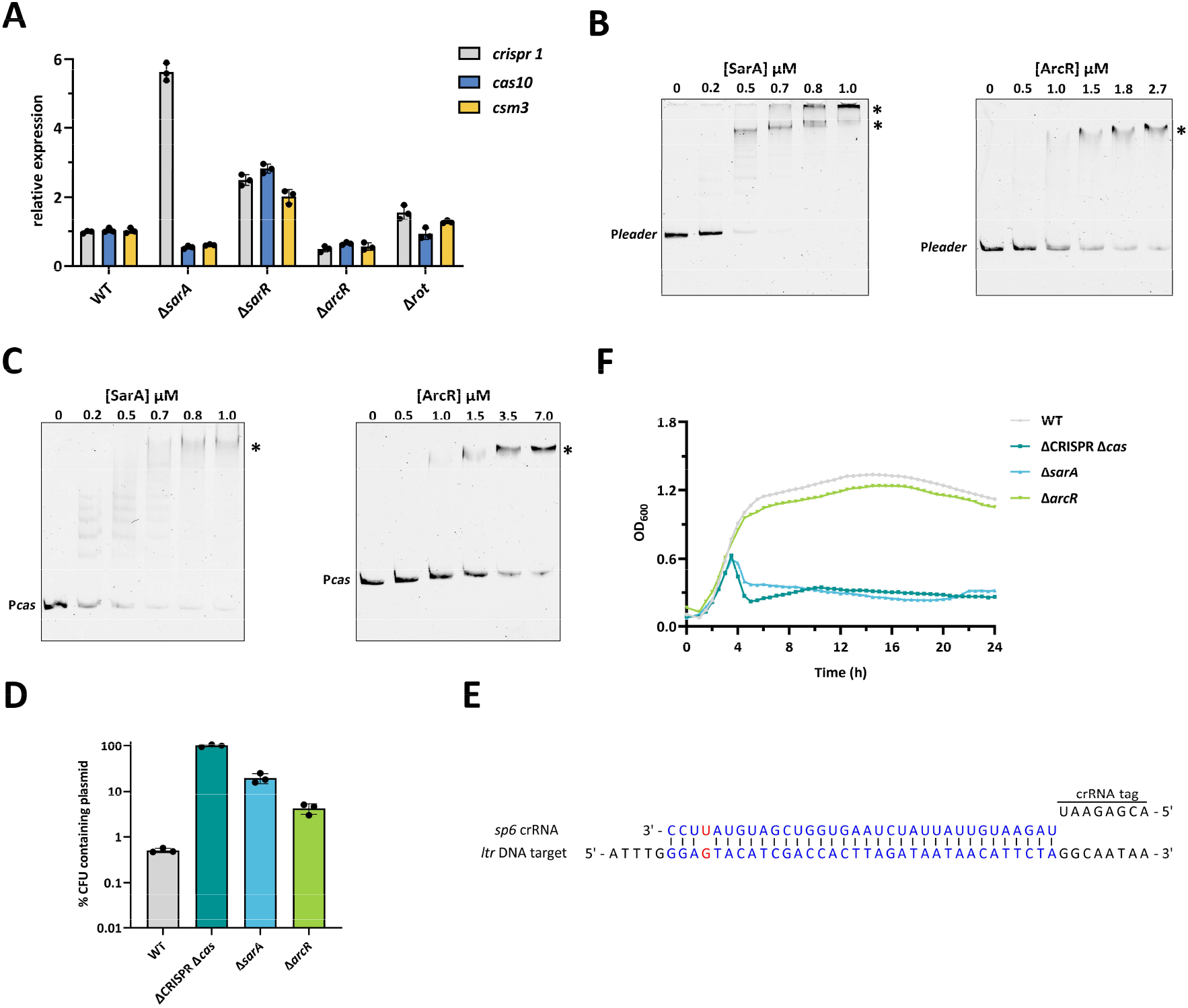
SarA and ArcR directly regulate the expression and activity of Type III-A CRISPR-Cas system. (A) qRT-PCR analysis of *cas10* and *csm3* expression in the WT and the individual mutant strains after 10 h of growth in liquid culture, respectively. (B) EMSAs were performed to analyze SarA and ArcR binding to the FAM-5’-end-labelled P*leader* sequence. DNA-protein complexes are indicated by an asterisk. (C) Same assay as in (B), using the FAM-5’-end-labelled P*cas* sequence. (D) Retention of the targeted plasmid pCR1SP1 in the WT, ΔCRISPR Δ*cas*, Δ*sarA* and Δ*arcR* mutants. The strains containing the plasmid were grown in TSB with ATc for 10 h followed by plating on TSA with or without chloramphenicol. % CFUs containing plasmid was scored as the number of Cm^R^ CFU/the total number of CFU. (E) Spacer 6 in the CRISPR1 array encodes a crRNA that targets the *ltr* gene of the *Staphylococcus* phage phiIPLA-RODI. There is one base (in red) mismatch between the *sp6* crRNA and the *ltr* protospacer. (F) Growth curve of the WT, ΔCRISPR Δ*cas*, Δ*sarA*, and Δ*arcR* mutants were infected with phage phiIPLA-RODI at an MOI of 0.1. All the data shown above (A, D and F) are means ± standard deviation of three independent experiments.

Since SarA, SarR, and ArcR affected the expression of *crispr1* and *cas* genes, we performed EMSA to test whether these regulators could directly bind to the P*leader* and P*cas* sequences, respectively. None of the three proteins shifted the promoter sequence of *gmk*, which served as a negative control (Figure S2B). As shown in Figure 2B and 2C, both SarA-His_6_ and ArcR-His_6_ proteins specifically shifted the P*leader* and P*cas* sequences, but SarR-His_6_ did not (Figure S2C). The specific binding of SarA-His_6_ and ArcR-His_6_ to the P*leader* and P*cas* sequences was confirmed by competitive EMSAs (Figure S2D and S2E). Collectively, these results demonstrate that SarA binds to the P*cas* promoter and activates *cas* genes expression but represses the transcription of CRISPR1 by interacting with the P*leader* sequence, and that ArcR functions as a direct positive regulator of both promoters.

### SarA and ArcR activate Type III-A CRISPR-Cas activity

The DNA cleavage activity of Cas10 is required for the Type III-A CRISPR-Cas immunity against phage infection.^12^ Therefore, we examined the effect of altered *cas10* expression on CRISPR-mediated immunity against the targeted plasmids and phages. To determine the effect of SarA and ArcR on CRISPR-Cas activity in eliminating invading foreign genetic elements, we measured the retention of an untargeted plasmid pRMC2 and a CRISPR-targeted plasmid pCR1SP1 in the WT, ΔCRISPR Δ*cas*, Δ*sarA*, and Δ*arcR* strains. The pCR1SP1 has a protospacer complementary to spacer 1 (*sp1*) in the CRISPR 1 locus of TZ0912. We used anhydrotetracycline (ATc) to induce the Pxy/tet-inducible promoter that drives transcription of the plasmid protospacer. The untargeted plasmid was completely retained in all the strains (Figure S2F). In contrast, we observed a robust clearance of the pCR1SP1 plasmid in the WT, confirming that the WT had effective CRISPR-Cas activity against the targeted plasmid. Compared with the retention of the pCR1SP1 plasmid in the WT, we observed 40-fold and 8-fold increase in plasmid retention in the Δ*sarA* and Δ*arcR* strains, respectively (Figure 2D).

Next, to determine whether SarA and ArcR could also regulate the Type III-A CRISPR-Cas system to defend against phage attacks, we exposed the mutant strains to phage phiIPLA-RODI, which is targeted by spacer 6 (*sp6*) at the CRISPR 1 locus of WT (Figure 2E), and monitored the bacterial growth over time in liquid culture. Compared to the WT strain, cells lacking the *sarA* gene succumbed to phage phiIPLA-RODI infection, similar to the ΔCRISPR Δ*cas* mutant (Figure 2F). However, the Δ*arcR* mutant exhibited a slight, but not significant, reduction in the immunity level relative to the WT (Figure 2F). These data demonstrate that SarA and ArcR can activate the interference of Type III-A CRISPR-Cas system.

### The binding sites for SarA and ArcR in P*leader* and P*cas* sequence

To further determine the binding sites of SarA and ArcR in P*leader* and P*cas* sequences, we divided the P*leader* and P*cas* sequence into seven and five ∼100 bp DNA fragments, respectively. The truncated DNA fragments were named according to the 5’ and 3’ positions of the P*leader* and P*cas* with respect to their transcriptional start site. There was a 50 bp overlap between these consecutive DNA fragments (Figure 3A and 3B). The binding between the DNA fragments and purified SarA-His_6_ or ArcR-His_6_ proteins was analyzed using EMSAs. We observed that ArcR-His_6_ specifically shifted the DNA fragments P*leader*-76+24 and P*leader*-26+74, but did not shift the DNA fragments P*leader*- 276-177, P*leader*-266-127, P*leader*-176-77, P*leader*-126-27, and P*leader*+25+112 (Figure 3C), indicating that the binding site for ArcR was located in P*leader*-26 and P*leader*+24. Binding of ArcR- His_6_ to the fragment P*leader*-26+24 was then confirmed by EMSA (Figure 3D). ArcR-His_6_ shifted the DNA fragment P*cas*-191-92, but not the other truncated P*cas* sequences (Figure 3E). The non-binding of ArcR-His_6_ to the DNA fragment P*cas*-241-142 and P*cas*-141-42 indicated that the ArcR-binding site in P*cas* was close to position P*cas*-141. Therefore, we generated three DNA fragments including position P*cas*-141 for EMSAs (Figure 3F). The results showed that both P*cas-*156-127 and P*cas*-161- 122 interacted with ArcR-His_6_, but not P*cas*-151-132 (Figure 3G). Previous studies revealed that ArcR in *S. aureus* was predicted to be a CRP family transcriptional regulator,^30^ and the conserved DNA binding motif of CRP is TGTGA-N_6_-TCACA in *E. coli*.^31^ Therefore, we examined whether the P*leader*-26+24 and P*cas-*156-127 fragments contain a conserved DNA-binding motif for ArcR. The homology analysis showed that the putative ArcR binding site is located between position +4 and +20 (5’-**TGTCA**AAAAAAG**TGACA**-3’) in the P*leader*-26+24 sequence and between position -152 and -137 (5’-**TGTGA**AGTTAT**TAGTA**-3’) in the P*cas-*156-127 sequence. However, the EMSA results showed that ArcR-His_6_ could not shift these two putative binding sites (Figure S3A), implying that the nucleotides flanking the putative binding site were also necessary for ArcR binding. Our results indicate that ArcR can specifically bind to P*leader*-26+24 and P*cas-*156-127 sequences (Figure S3B and S3C).

**Figure 3.**
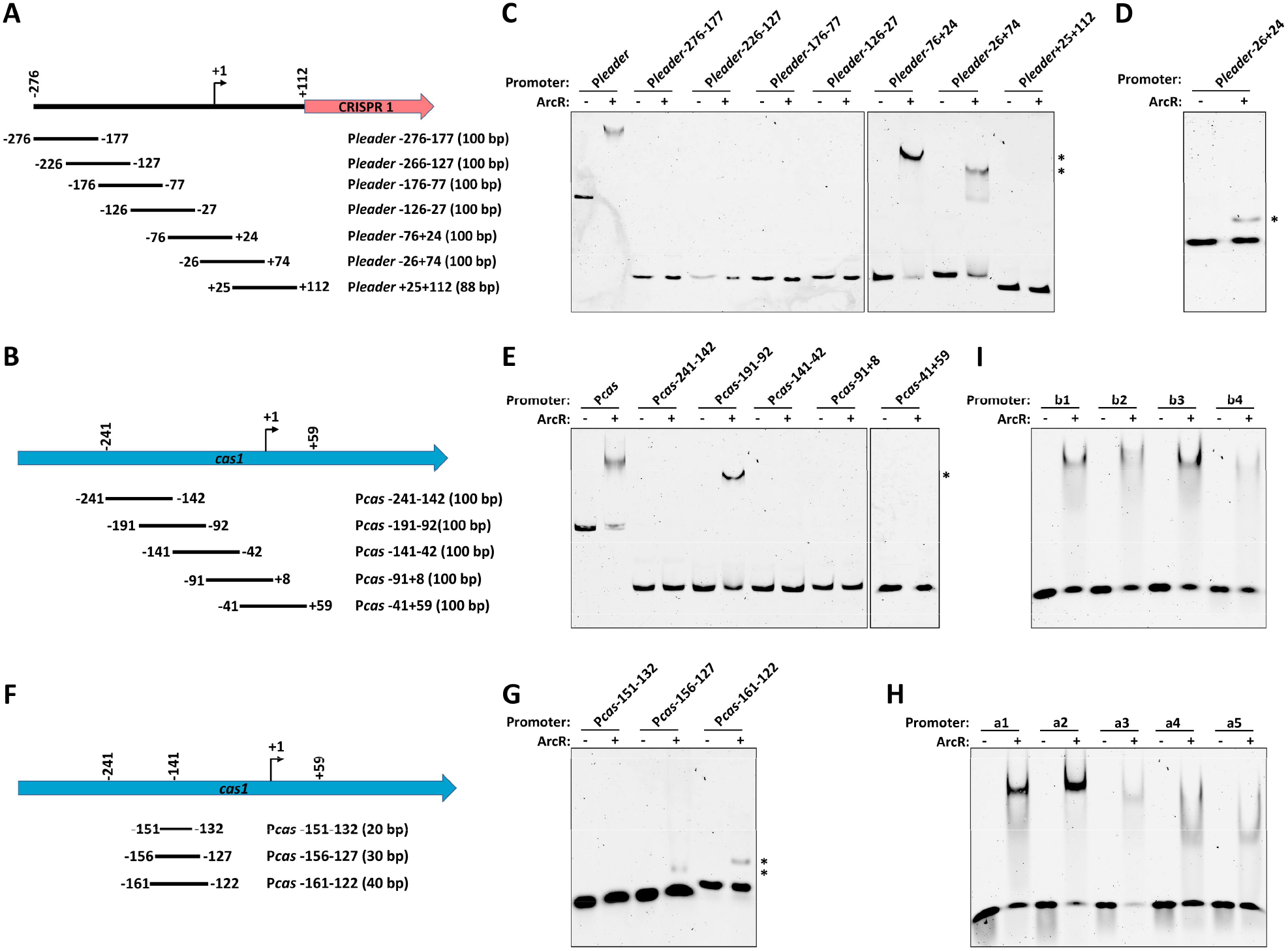
Binding sites for SarA and ArcR in the P*leader* and P*cas* sequences. (A and B) Schematic representation of P*leader* (A) and P*cas* (B) loci. Various ∼50 bp overlapping DNA fragments covering different regions of P*leader* and P*cas* were tested in EMSA experiments. The transcriptional start sites (+1) of P*leader* and P*cas* are indicated by bent arrows. All positions indicated are relative to the transcriptional start site. (C) Seven truncated FAM-5’-end-labelled P*leader* fragments were incubated with ArcR-His_6_ at a concentration of 7.2 μM. (D) FAM-5’-end-labelled P*leader*-26+24 fragment was incubated with 7.2 μM ArcR-His_6_. (E) Five truncated FAM-5’-end-labelled P*cas* fragments were incubated with 7.2 μM ArcR-His_6_. (F) Schematic representation of three DNA fragments flanking position -141 of P*cas*. (G) Three FAM-5’-end-labelled DNA fragments were incubated with ArcR-His_6_ at the concentration of 7.2 μM, respectively. (H and I) The FAM-5’-end-labelled putative SarA-binding sites on P*leader* (a1-a5) (H) and P*cas* (b1-b4) (I) fragments were incubated with SarA-His_6_ at a concentration of 1.0 μM, respectively. DNA-protein complexes are indicated by asterisks. +, with protein; –, without protein.

Moreover, EMSAs showed that the SarA-His_6_ could bind to each truncated P*leader* and P*cas* sequence (Figure S3D and S3E), implying that multiple SarA-binding sites were present in the P*leader* and P*cas* sequences. SarA has been reported to have a preference for AT-rich binding sites ^32,33^, and the consensus SarA-binding motif has been predicted to be ATTTTAT ^32,34^. In agreement, we identified five and four putative SarA-binding sites in the P*leader* (a1-a5) and P*cas* (b1-b4) sequences, respectively (Figure S3B and S3C). EMSAs confirmed that SarA-His_6_ could bind to a1-a5 and b1-b4 binding sites (Figure 3H and 3I). Notably, we found that the a4 and b2 SarA-binding sites in the P*leader* and P*cas* sequences were located close to ArcR-binding sites (Figure S3B and S3C), indicating that SarA and ArcR might compete for interactions with the P*leader* and P*cas* sequences.

### SarA and ArcR compete for binding to the P*leader* and P*cas* sequence

Our above results showed that SarA and ArcR binding sites were located close to each other, indicating a potential competition between SarA and ArcR. Thus, we performed competitive EMSAs to investigate if SarA and ArcR could compete for binding to the P*leader* and P*cas* sequences. The P*leader* or P*cas* fragment was preincubated with ArcR at room temperature for 30 min, followed by adding increasing concentrations of SarA. As shown in Figure 4A and 4B, the increasing concentrations of SarA shifted the P*leader*-ArcR or P*cas*-ArcR complex to a slower-migrating complex similar to that formed by SarA alone, suggesting that SarA could displace ArcR from the P*leader* and P*cas* sequences. A similar assay was performed by first incubating the P*leader* and P*cas* fragments with SarA, followed by adding increasing concentrations of ArcR. The results showed that ArcR could also displace SarA from the P*leader* and P*cas* sequence. However, this displacement of SarA required a much higher ArcR concentration of > 3.6 μM (Figure 4C and 4D). Together, our data demonstrates that SraA is more competitive than ArcR in term of interaction with P*leader* and P*cas* sequences.

**Figure 4.**
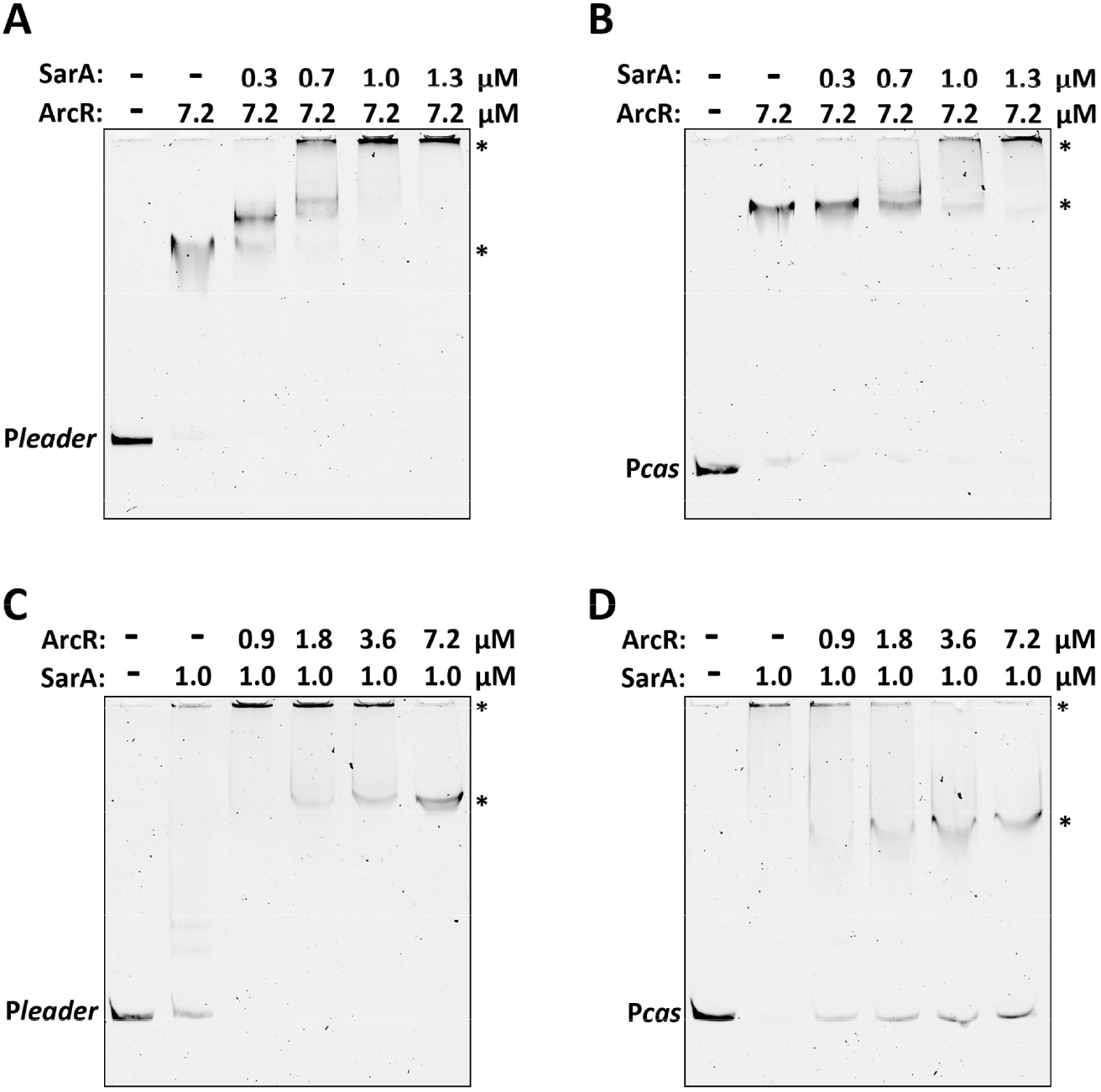
Competition between SarA and ArcR binding to P*leader* and P*cas* sequences using competitive EMSAs. (A and B) FAM-5’-end-labelled P*leader* sequence (A) or P*cas* sequence (B) was first incubated with 7.2 μM purified ArcR-His_6_ followed by the addition of increasing concentrations of SarA-His_6_ (from 0.3 μM to 1.3 μM). The shifted bands are indicated by arrows. (C and D) FAM-5’- end-labelled P*leader* sequence (C) or P*cas* sequence (D) was first incubated with 1.0 μM purified SarA-His_6_ followed by the addition of increasing concentrations of ArcR-His_6_ (from 0.9 μM to 7.2 μM). The shifted bands are indicated by arrows.

### Quorum Sensing indirectly represses expression of *cas* genes in the Type III-A CRISPR-Cas system

In *S. aureus*, SarA is a cytoplasmic transcriptional regulator promoting expression the QS system, thereby enhancing virulence factors expression.^35^ ArcR is a member of the CRP-family transcriptional regulators, and it can activate the expression of *arcABDC* genes, which is responsible for the utilization of arginine as an energy source for growth under anaerobic conditions.^30^ Both transcriptional regulators are reported to be related to QS regulation,^35,36^ indicating that QS may have an effect on regulation of the CRISPR-Cas system in *S. aureus*. Although QS-mediated activation of CRISPR-Cas systems have been reported in several gram-negative bacteria, the impact of QS in CRISPR-Cas system remains poorly understood in gram-positive bacteria. To characterize the growth phase of the TZ0912 strain at low cell density and high cell density, we monitored CFU counts and *agrA* expression levels for QS signal every two hours. At 2 h, the cell density was 3.63×10^8^ CFU/mL with a low *agrA* expression level. The expression of *agrA* reached a peak at 10 h with a cell density of 5.20×10^9^ CFU/mL, while *agrA* expression decreased after 10 h (Figure 5A). Therefore, we used 2 h and 10 h to represent the low cell density and high cell density, respectively. Subsequently, we measured the expression of *cas10* and *csm3* at both the low cell density and high cell density using qRT-PCR. Both *cas10* and *csm3* genes were highly expressed at low cell density, while their expression was decreased at high cell density (Figure S4A and S4B), suggesting that the expression of these genes may be repressed in response to QS signaling at high cell density. To determine the effect of QS on CRISPR-*cas* expression, we constructed a Δ*agr* mutant by inactivating the *agr*BDCA operon and the RNA III gene, and subsequently quantified the *cas* genes expression in the wild type (WT) and Δ*agr* mutant strains. At low cell density, the expression of both *cas* genes was similar in WT and Δ*agr* mutant cells, whereas at high cell density, expression of both *cas10* and *csm3* was significantly higher in the Δ*agr* mutant compared to the expression levels in the WT (Figure S4A and S4B).

**Figure 5.**
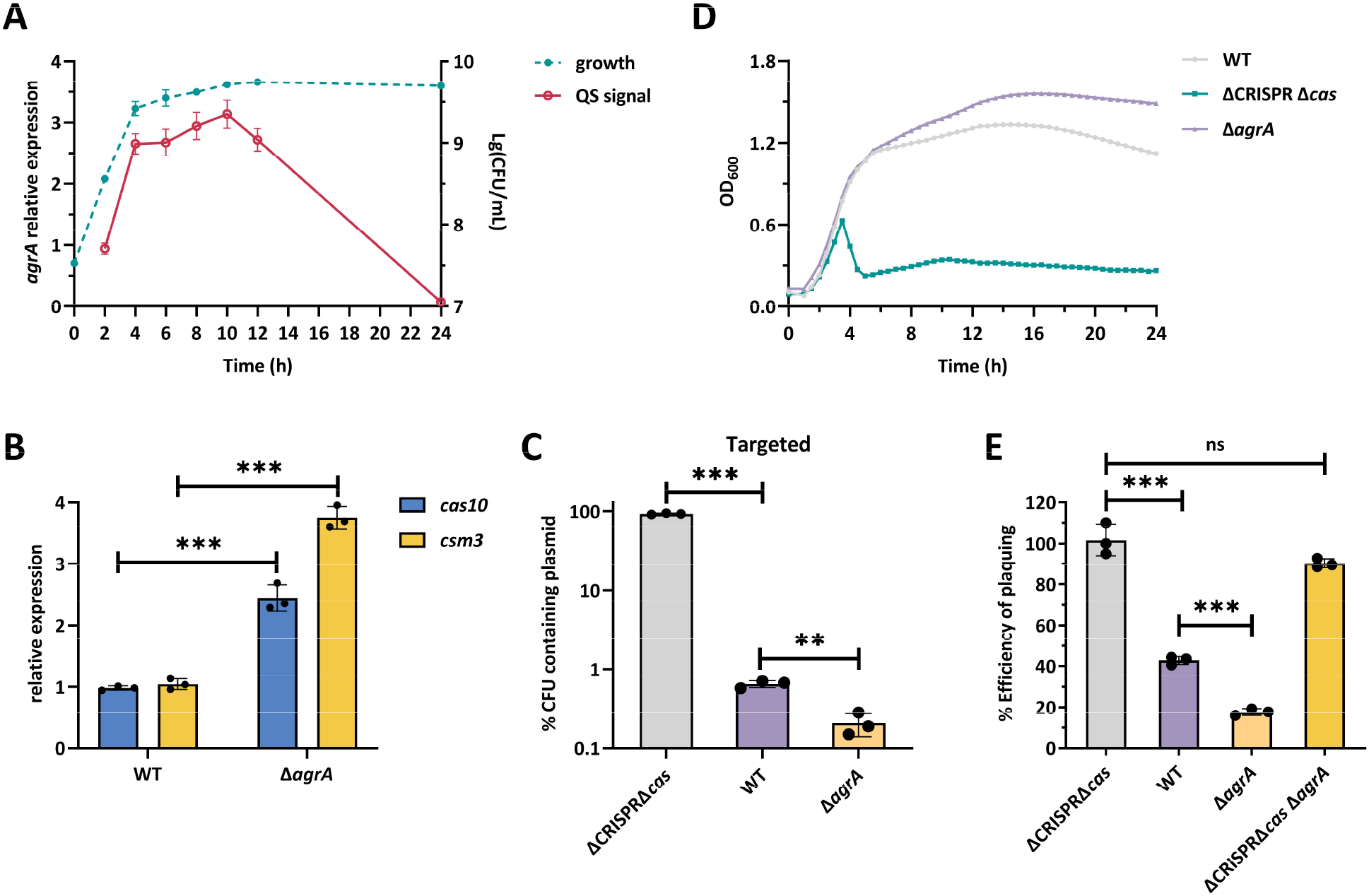
QS represses the expression and interference of Type III-A CRISPR-Cas system in *S. aureus*. (A) Growth curve of *S. aureus* TZ0912 strain and *agrA* mRNA levels. The transcriptional level of *agrA* was measured by qRT-PCR. The *gyrB* was used as an internal control. (B) qRT-PCR analysis of *cas10* and *csm3* expression in the WT and Δ*agrA* mutant at high cell density. (C) Retention of the targeted plasmid pCR1SP1 in the WT, ΔCRISPR Δ*cas* and Δ*agrA* mutant. The strains containing the plasmid were grown in TSB with ATc for 10 h followed by plating on TSA with or without chloramphenicol. % CFUs containing plasmid was scored as the number of Cm^R^ CFU/the total number of CFU. (D) Growth curve of the WT, ΔCRISPR Δ*cas*, and Δ*agrA* mutant were infected with phage phiIPLA-RODI at an MOI of 0.1. (E) The immunity against phage phiIPLA-RODI of relevant strains was quantified by efficiency of plaquing, normalized to the number of PFU/mL measured on the ΔCRISPR Δ*cas* mutant. All data (B-E) are means ± standard deviation of three independent experiments. A two-tailed unpaired Student’s *t*-test was used to calculate *P* values; \*\**p*<0.01, \*\*\**p*<0.001, ns, not significant.

Subsequently, we analyzed the effects of AgrA and RNA III on CRISPR-*cas* expression in *S. aureus*. Compared with the WT, the deletion of *RNA III* led to an ∼1.8-fold increase in *crispr1* expression, but no effect was observed on *cas* genes and *crispr2* expression (Figure S4C). However, the inactivation of *agrA* in WT increased the expression of *cas* genes to the same level as that in the Δ*agr* mutant at high cell density (Figure 5B), indicating that AgrA negatively regulated *cas* genes expression in *S. aureus*. Furthermore, the *crispr1* expression was significantly higher in the Δ*agrA* mutant than in the WT (2.5-fold), whereas the expression of *crispr2* in the Δ*agrA* mutant was not significantly affected (Figure S4D). These data suggest that QS regulation of the Type III-A CRISPR-Cas system is mediated by AgrA, which functions as a repressor of *cas* genes.

To determine whether AgrA directly interacts with the promoter regions, we examined the interaction of AgrA with the CRISPR-Cas P*leader* and P*cas* sequences using EMSAs. As shown in Figure S2E, AgrA-His_6_ protein shifted the P2 promoter of the *agr* locus, starting at a concentration of 0.25 μM, as expected,^37^ showing that the tagged protein was functional. However, the P*leader* and P*cas* DNA fragments did not shift in the presence of AgrA-His_6_ up to a concentration of 2.5 μM, indicating that AgrA could not bind specifically to these promoter regions. Our aforementioned results suggested that AgrA was not captured in the DNA pull-down assays, further indicating that AgrA represses the expression of *cas* genes indirectly and that other regulatory factors downstream of AgrA may underlie the QS-mediated repression of *cas* genes.

### Quorum Sensing regulates Type III-A CRISPR-Cas activity

To determine the effect of QS on CRISPR-Cas activity in eliminating invading foreign genetic elements, we measured the retention of an untargeted plasmid pRMC2 and a CRISPR-targeted plasmid pCR1SP1 in the WT, ΔCRISPR Δ*cas*, and Δ*agrA* strains. As expected, the untargeted plasmid was completely retained in all strains (Figure S4F). We observed that the Δ*agrA* mutant showed a 3-fold decrease in the retention of the pCR1SP1 plasmid compared to the WT (Figure 5C), indicating that AgrA represses the interference activity of the Type III-A CRISPR-Cas system against the plasmid.

To examine the effect of QS on CRISPR-Cas activity against phage infections, we exposed the WT, ΔCRISPR Δ*cas*, and Δ*agrA* strains to phage philPLA-RODI. Consistent with the increased *cas10* expression in Δ*agrA* mutant, the Δ*agrA* mutant was more resistant to phage philPLA-RODI than the WT (Figure 5D). A similar result was obtained when we measured the efficiency of plaquing (EOP) of phiIPLA-RODI on these strains (Figure 5E). We observed that the CRISPR-targeted phage philPLA-RODI showed an EOP of 40% in the WT strain compared to the ΔCRISPR Δ*cas* mutant, whereas an EOP of 17% was observed in the Δ*agrA* mutant relative to the ΔCRISPR Δ*cas* mutant (Figure 5E).

To eliminate other QS-regulated factors that may confound our analysis, e.g. any effects of QS on the expression of the phage receptor, as observed for other phage-host pairs,^38^ we deleted *agrA* in the ΔCRISPR Δ*cas* mutant and infected the triple mutant with phage philPLA-RODI. The *Staphylococcus* phage phiSA012, without homology to any spacers in WT, was used as a non-CRISPR targeted control. Both phages showed similar EOP on the ΔCRISPR Δ*cas* Δ*agrA* and ΔCRISPR Δ*cas* mutants (Figure 5E and S4G), indicating that QS does not affect the ability of the phages to infect this host. Collectively, these results demonstrate that QS represses Type III-A CRISPR-Cas interference in *S. aureus*.

### AgrA represses Type III-A CRISPR-Cas expression and interference through downregulation of SarA and ArcR regulators

At high cell density, qRT-PCR analysis showed that both *sarA* and *arcR* were upregulated in the Δ*agrA* mutant compared to that in the WT (Figure 6A). EMSA further confirmed that AgrA directly repressed the expression of *sarA* and *arcR* by binding to their promoters. (Figure 6B and S5A). Therefore, we hypothesized that AgrA may repress Type III-A CRISPR-Cas expression by inhibiting SarA- and ArcR-mediated activation of *cas* genes. To confirm our hypothesis, we constructed the double mutants Δ*agrA* Δ*sarA*, Δ*agrA* Δ*arcR*, and Δ*sarA* Δ*arcR* and a triple mutant Δ*agrA* Δ*sarA* Δ*arcR*, and quantified the expression of *cas10* by qRT-PCR. Deletion of *arcR* or *sarA* restored *cas10* expression in the Δ*agrA* mutant to the WT level (Figure 6C). Importantly, the level of *cas10* gene expression was equivalent between the Δ*sarA* Δ*arcR* mutant and the Δ*agrA* Δ*sarA* Δ*arcR* mutant (Figure 6C), indicating that AgrA represses *cas* gene expression in a SarA/ArcR-dependent manner. These results indicate that when the bacteria grow to a high cell density, AgrA represses CRISPR-Cas expression via inhibiting the activation of *cas* genes by SarA and ArcR.

**Figure 6.**
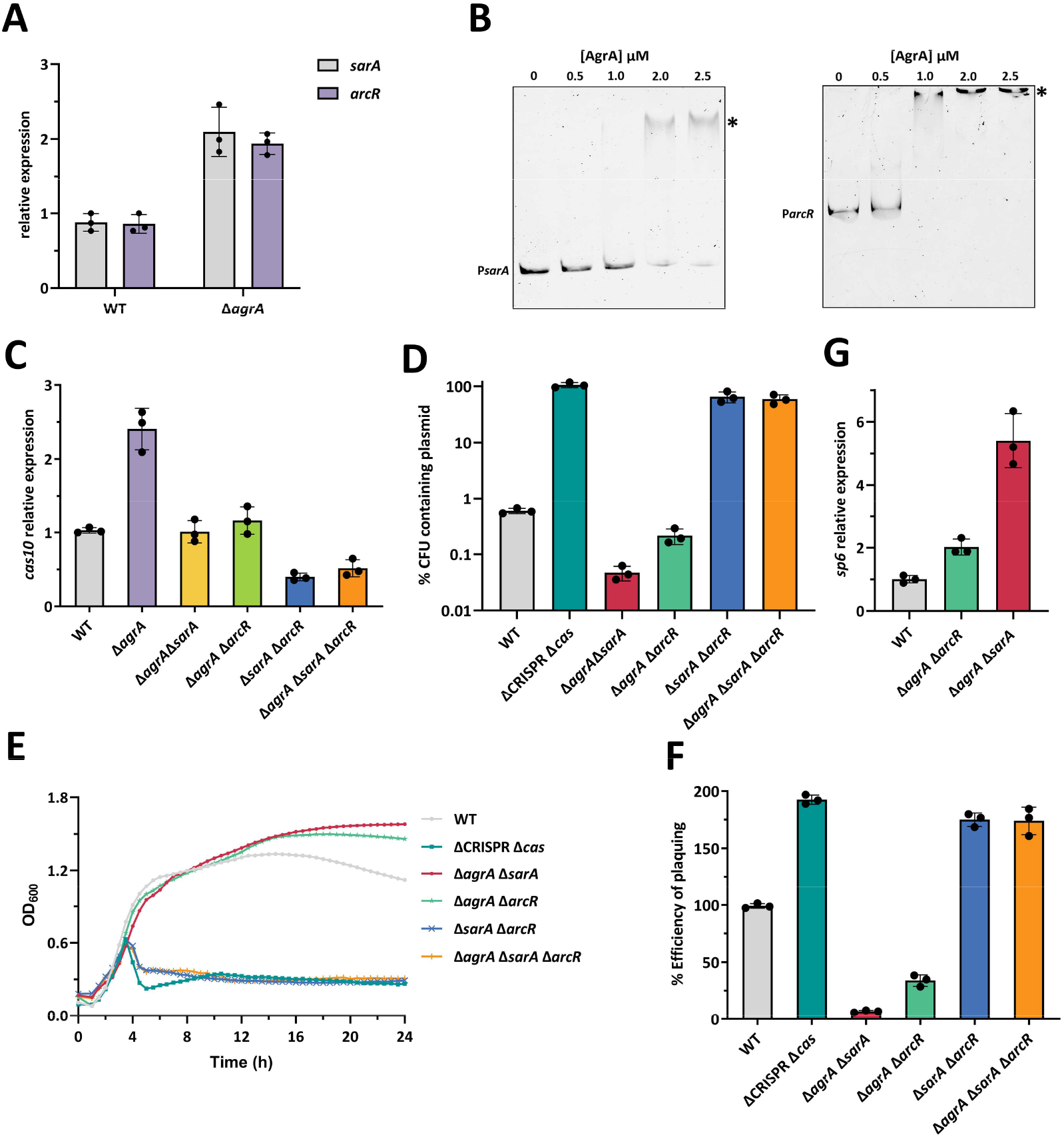
The AgrA-mediated regulation of CRISPR-Cas is dependent on SarA and ArcR. (A) qRT-PCR measurement of *sarA* and *arcR* expression in the WT and Δ*agrA* mutant after 10 h of growth in liquid culture, respectively. (B) FAM-5’-end-labelled P*sarA* sequence and P*arcR* sequence was incubated with the increasing concentrations of purified AgrA-His_6_ protein. Acetyl phosphate (50 mM) was added to all EMSAs. DNA-protein complexes are indicated by an asterisk. (C) qRT-PCR measurement of *cas10* expression in the WT, Δ*agrA*, and double mutants (Δ*agrA* Δ*sarA*, Δ*agrA* Δ*arcR*, Δ*sarA* Δ*arcR*), and triple mutant Δ*agrA* Δ*sarA* Δ*arcR* after 10 h of growth in liquid culture, respectively. (D) Retention of the targeted plasmid pCR1SP1 in the WT, ΔCRISPR Δ*cas*, and double mutants (Δ*agrA* Δ*sarA*, Δ*agrA* Δ*arcR*, Δ*sarA* Δ*arcR*), and triple mutant Δ*agrA* Δ*sarA* Δ*arcR*. The strains containing the pCR1SP1 plasmid were grown in TSB with ATc for 10 h followed by plating on TSA with or without chloramphenicol. % CFUs represents the ratio of bacteria grown on plates containing chloramphenicol to the number of bacteria growing on plates without antibiotics. (E) Growth curve of the WT, ΔCRISPR Δ*cas*, and double mutants (Δ*agrA* Δ*sarA*, Δ*agrA* Δ*arcR*, Δ*sarA* Δ*arcR*), and triple mutant Δ*agrA* Δ*sarA* Δ*arcR* were infected with phage phiIPLA-RODI at an MOI of 0.1. (F) The immunity against phage phiIPLA-RODI of relevant strains was represented as efficiency of plaquing, which is a ratio relative to the number of plaques measured on the WT. (G) qRT-PCR measurement of *spc6* expression in the WT, and double mutants (Δ*agrA* Δ*sarA*, Δ*agrA* Δ*arcR*) after 10 h of growth in liquid culture, respectively. All the data (A, C-F) are means ± standard deviation of three independent experiments.

Next, we investigated whether the inhibitory effect of AgrA on the activity of the Type III-A CRISPR-Cas system was also mediated through SarA and ArcR. Interference against plasmids was assessed by measuring the retention of the targeted plasmid pCR1SP1 and the untargeted plasmid in the WT, Δ*agrA* Δ*sarA*, Δ*agrA* Δ*arcR*, Δ*sarA* Δ*arcR* and Δ*agrA* Δ*sarA* Δ*arcR* mutant strains. As expected, none of the strains lost the untargeted plasmid (Figure S5B). Compared with the retention of the pCR1SP1 plasmid in the WT, the deletion of both *sarA* and *arcR* in the Δ*agrA* mutant abolished Type III-A against the targeted plasmid, which was also observed in the Δ*sarA* Δ*arcR* mutant (Figure 6D), indicating that AgrA-mediated repression was dependent on SarA and ArcR. Besides, a decrease in pCRISP1 retention was observed in the Δ*agrA* Δ*sarA* and Δ*agrA* Δ*arcR* mutants, indicating that both activated ArcR and SarA could promote CRISPR-Cas expression and interference activity (Figure 6D).

Moreover, we exposed the mutant strains to phiIPLA-RODI and monitored the bacterial growth over time in liquid culture. Compared to the WT strain, the Δ*sarA* Δ*arcR* and Δ*agrA* Δ*sarA* Δ*arcR* mutants failed to provide immunity (Figure 6E), consistent with the reduced *cas10* expression. Although *cas10* expression in the Δ*agrA* Δ*sarA* and Δ*agrA* Δ*arcR* mutants was restored to the level of the WT, these two mutants showed more effective antiphage immunity than the WT (Figure 6E). A similar result was obtained when we measured the EOP of phiIPLA-RODI on these mutant strains (Figure 6F). We have previously demonstrated that the expression level of crRNA is also closely related with CRISPR-Cas immune efficiency in the WT,^20^ which has also been shown in *Streptococcus pyogenes* (*S. pyogenes*) Type II-A CRISPR-Cas system.^39^ Therefore, we analyzed the transcriptional level of *sp6* in the WT, Δ*agrA* Δ*arcR*, and Δ*agrA* Δ*sarA* strains and found increased *sp6* expression levels in both mutant strains compared to the WT strain (Figure 6G), indicating that the increased antiphage immunity against phiIPLA-RODI in both mutant strains was due to higher expression of *sp6* than that in the WT strain. These results demonstrate that AgrA can repress the Type III-A CRISPR-Cas interference in *S. aureus* TZ0912 by inhibiting both SarA and ArcR, which are positive regulators of the Type III-A CRISPR-Cas system.

### P*leader* and P*cas* sequence are conserved in staphylococcal Type III-A CRISPR-Cas system

To investigate whether the identified P*leader* and P*cas* sequence is conserved in Firmicutes, we performed BLAST analysis against the NCBI nr nucleotide database limited to Firmicutes, using the identified P*leader* and P*cas* sequence as queries. A comprehensive list of P*leader* and P*cas* homologs was listed in Table S3 and Table S4, respectively. P*leader* homologs are mainly distributed in *Staphylococcus* of Firmicutes (Table S3). Maximum likelihood phylogenetic analysis of the P*leader* homologs revealed that most homologs are highly conserved in *Staphylococci* carrying the Type III-A CRISPR-Cas system (Figure S6A). In comparison, the P*cas* sequence is widespread in various Firmicutes, including *Staphylococcaceae, Listeriaceae, Aerococcaceae, Lactobacillaceae, Enterococcaceae* and *Streptococcaceae* family (Table S4). The phylogenetic tree of the P*cas* homologs was divided into three clades, which correspond to the related bacterial families (Figure S6B). The high conservation of P*cas* sequences in the Type II and III CRISPR-Cas system in Firmicutes indicated that P*cas* was a conserved promoter of *cas* genes in both CRISPR-Cas systems (Figure S6B). All P*cas* sequence from staphylococcal Type III-A CRISPR-Cas system was conserved, which clustered together in the phylogenetic tree. Meanwhile, the occurrence of the QS system was also examined in the P*leader*- and P*cas*-positive strains. We found that all identified P*leader*- and P*cas*-positive strains encoded a QS system (Figure S6A and S6B). Interestingly, the *comABCDE* operon in *Streptococcus* and *fsrABDC* operon in *Enterococcus* encode a *S. aureus agr*-like two-component regulatory system,^40,41^ suggesting that *agr*-mediated repression of the CRISPR-Cas system may be common phenomenon in Firmicutes. Altogether, these observations suggest that the P*leader* and P*cas* sequence are conserved in staphylococcal Type III-A CRISPR-Cas system.

## DISCUSSION

Although the structure and immunity mechanism of the Type III-A CRISPR-Cas systems in *Staphylococci* have been studied extensively, the regulatory pathways governing the expression of this system are underexplored. We showed that the CRISPR array and *cas* genes are transcribed as a single transcript, whereas the *cas* genes downstream of *cas1* can also be transcribed using its own promoter, P*cas*, located within *cas1*. The presence of a promoter in the CRISPR leader sequence is in agreement with the Type I CRISPR-Cas systems in *E. coli* and *Pectobacterium atrosepticum* (*P. atrosepticum*) also harbors promoters in their leader sequences.^42,43^ In addition to its ability to initiate crRNA transcription, the leader sequence can also direct the integration of new spacers into the first repeat in the Type II-A CRISPR-Cas system.^39,44^ In contrast to previous studies reporting that *cas1* has no effect on antiplasmid immunity in *S. aureus* and *S. epidermidis* Type III-A CRISPR-Cas system,^21,45^ our results showed that mutation of *cas1* led to weaker antiphage immunity than the WT strain, likely because this abolishes expression of the downstream *cas* genes. Our RNA-seq results showed that the transcriptional expression level of *cas1* gene was not abundant, which probably explains why spacer adaptation was not frequently observed in Type III-A CRISPR-Cas system under the experimental conditions.

Several transcriptional regulators have been shown to control the expression of CRISPR-*cas* systems.^46^ For instance, the histone-like nucleoid-structuring (H-NS) DNA-binding protein was reported to repress the Type I-E CRISPR-*cas* expression in *E. coli*.^43^ The extracytoplasmic function (ECF) σ factor DdvS has also been implicated in Type III-B CRISPR-*cas* activation by binding to the promoter of *cas* genes in *Myxococcus xanthus*.^47^ Likewise, we find that the regulation of Type III-A CRISPR-Cas system in *S. aureus* is controlled by transcriptional regulators SarA and ArcR. In *S. aureus*, SarA is a global transcriptional regulator that binds to an AT-rich site. It can modulate expression of multiple target genes involved in virulence.^48^ Mutation of *sarA* in *S. aureus* resulted in a significant decrease in biofilm formation, indicating that SarA is crucial for biofilm formation.^49^ Several studies have confirmed that phage therapy is an important method for inhibition of *S. aureus* biofilm formation.^50,51^ Here, we show that SarA functions as a positive regulator of the CRISPR-Cas system, thus preventing bacteria from phage attack. Our study connects the regulation of biofilm formation and the activity of CRISPR-Cas system to the common regulator SarA, suggesting that SarA-mediated activation of CRISPR-Cas system could act as an alternative anti-phage defense mechanism when biofilms are destroyed by phages. Additionally, our results indicated that the transcriptional regulator SarR can indirectly repress the expression of *cas* genes. SarR is a negative regulator of *sarA* expression in *S. aureus*.^52^ Therefore, it is likely that SarR decreases CRISPR-*cas* expression by repressing the direct regulator SarA.

ArcR can control the expression of numerous genes required for anaerobic growth of *S. aureus* by binding to a conserved CRP-like sequence motif TGTGA-N_6_-TCACA.^30,53^ We determined that ArcR activates the Type III-A CRISPR-Cas activity by binding to the CRP-like sequence motif in the P*leader* and P*cas* sequences, which are widespread in the Firmicutes CRISPR-Cas system. CRP-mediated regulation of Type I and III CRISPR-Cas systems has also been reported in many bacteria,^54-57^ suggesting that it might be a general phenomenon in diverse bacteria. ArcR utilizes arginine as an energy source for growth under anaerobic conditions, which can be repressed by glucose in *S. aureus*.^30^ Hence, the ArcR-activated CRISPR-Cas system can be suppressed when glucose serves as the main energy source, thereby reducing unnecessary resource consumption and providing more energy for bacterial growth, which is consistent with the findings observed in *P. atrosepticum*.^55^ Notably, deletion of *arcR* alone resulted in a slightly weaker immunity against plasmid and phage infection. However, it is interesting that deletion of *arcR* in the Δ*sarA* or Δ*agrA* Δ*sarA* mutant completely abolished the immunity of CRISPR-Cas system, which was similar to Δ*sarA* mutant, indicating that ArcR could provide a strong immunity in the absence of SarA. This could be explained by the fact that SarA was more competitive than ArcR in binding to the P*leader* and P*cas* fragments because several SarA- binding sites were present in the P*leader* and P*cas* sequences. Alternatively, since ArcR exerts its regulatory function under anaerobic conditions,^30^ it is possible that ArcR has a much less pronounced effect on immunity under the physiological culture conditions employed in this study. Therefore, future work will be required to completely illuminate the mechanism of ArcR on CRISPR-Cas immunity.

QS is a universal phenomenon observed across Gram-negative and Gram-positive bacteria, and it allows bacteria to coordinate biological functions in response to cell density and microbial composition.^24^ When the bacterial population reaches a high cell density, bacteria are at an increased risk of phage infection, because phages successfully spread in an environment with an abundance of susceptible hosts.^26^ Accordingly, bacteria use QS to activate different phage defense systems. For example, both *E. coli* and *Vibrio anguillarum* use QS to downregulate the expression of phage receptors to prevent phage infection.^38,58^ Similarly, the QS-mediated activation of CRISPR-Cas systems has been demonstrated in *P. aeruginosa* and *Serratia*.^22,23^ However, we observed the opposite role of QS in regulating the *S. aureus* Type III-A CRISPR-Cas system.

The repression of CRISPR-Cas could increase the susceptibility of bacteria to phage attacks. Therefore, other innate immune systems, including surface modifications, restriction modifications, and abortive infection systems, are required to interfere with phage infection. An abortive infection system in *Staphylococci* was shown to be activated by serine/threonine kinases, thereby preventing phage release and protecting the bacterial population.^59^ In *Serratia*, the Rcs membrane stress response pathway represses all three CRISPR-Cas systems (Type I-E, I-F and III-A) to promote the uptake of resistance genes, while it enhances the surface-based innate immunity against phage infection.^60^ In *S. pyogenes*, a natural single-guide RNA can direct Cas9 to repress the Type II-A CRISPR-Cas activation to avoid autoimmunity at the cost of reduced immunity against phages.^61^ Hence, tightly coordinated and balanced regulation of adaptive and innate immune systems would allow bacteria to optimize their gene acquisition and defense systems according to environmental needs. Since certain proteins encoded by bacteria can inhibit the expression and interference of CRISPR-Cas systems, plasmids or phages are likely to obtain these protein homologues in response to CRISPR-Cas immunity. Indeed, it was recently reported that homologues of the CRISPR-*cas* suppressors AmrZ and RsmA are widely present on plasmids or phages,^62,63^ implying that these proteins potentially function to overcome CRISPR-Cas immunity.

In summary, we found that the AgrA QS system could indirectly repress the expression and interference of Type III-A CRISPR-Cas system in *S. aureus*. We identified two transcriptional regulators, SarA and ArcR, that could activate the *cas* genes expression by binding directly to the CRISPR1 and *cas* promoters. Importantly, we further demonstrated that AgrA could inhibit the expression of *sarA* and *arcR*, underpinning the AgrA-mediated regulation of the CRISPR-Cas system in *S. aureus* (Figure 7).

**Figure 7.**
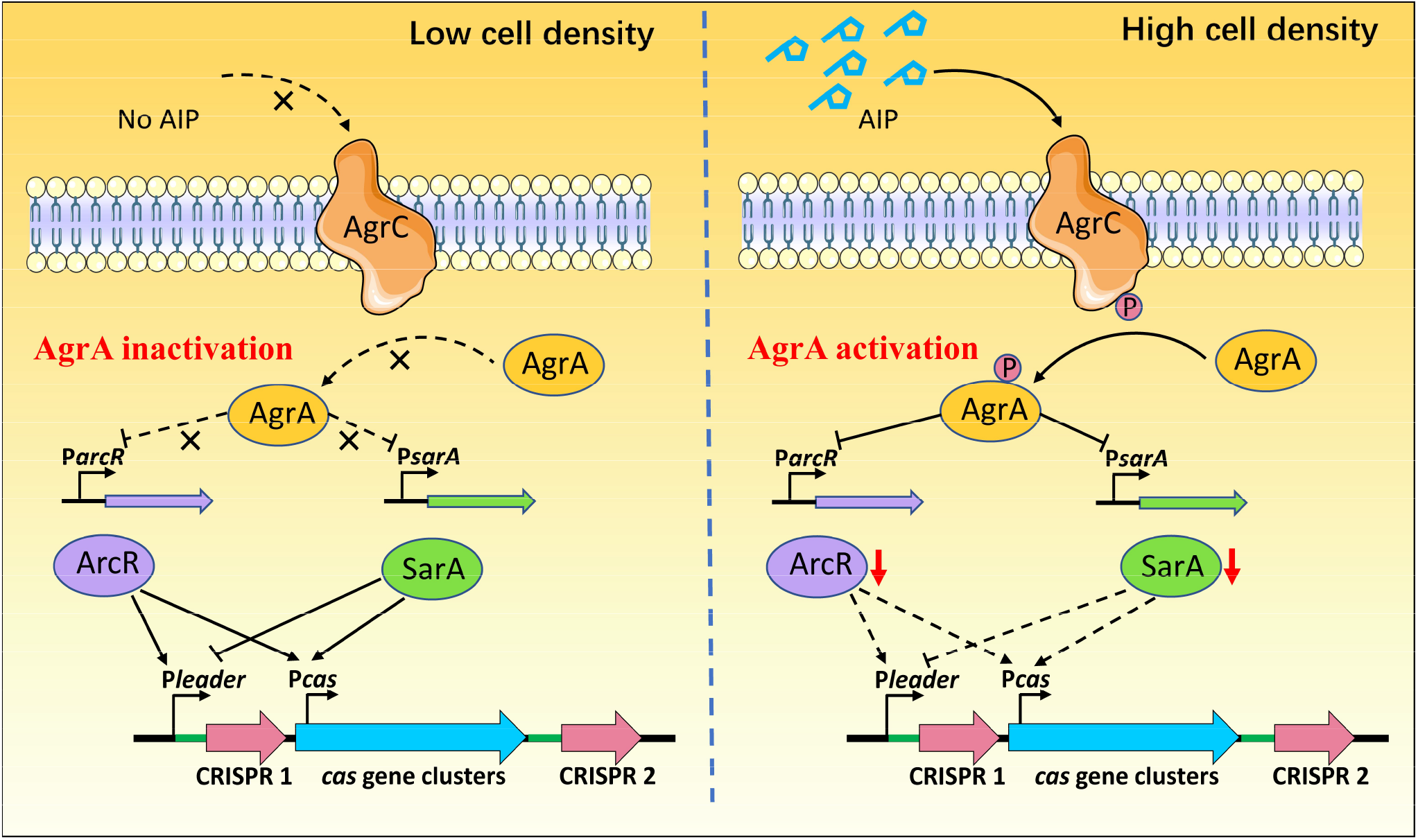
Model of QS-mediated regulation of the Type III-A CRISPR-Cas system in *S. aureus*. At low cell density, no AIP are sensed by AgrC, which cannot phosphorylate and thus activate AgrA. SarA and ArcR activate *cas* gene expression by binding to the P*cas* sequence. However, when the bacteria grow to high cell density, the AIP concentration reaches a threshold and activate the kinase domain of AgrC, enabling autophosphorylation. AgrC then phosphorylates and activates AgrA. The activated AgrA in turn represses the expression of SarA and ArcR, thereby inhibiting the direct activation of *cas* by SarA and ArcR.

## ACKNOWLEDGEMENTS

We thank Dr. Pilar García for providing the *Staphylococcus* phage phiIPLA-RODI and Dr. Xiancai Rao for providing the plasmid pOS1-*lacZ*. We thank Dr. Dan Gu for helpful guidance in EMSA and DNA-pull down assays. This study was supported by Natural Science Foundation of Jiangsu Province (BK20190883); The Natural Science Foundation of the Jiangsu Higher Education Institutions of China (19KJB230007); the fifth phase of “333 project” scientific research project in Jiangsu Province (BRA2020002); National Natural Science Foundation of China (32072821, 31730094); The Priority Academic Program Development of Jiangsu Higher Education Institution (PAPD); The Postgraduate Research & Practice Innovation Program of Jiangsu Province (KYCX21_3267); NMHK was supported by the Lundbeck Foundation Grant R264-2017-3936.

## AUTHOR CONTRIBUTIONS

L.Q.C., J.X.A., H.I., N.M.H.K. and L.Y. conceived and designed the study; L.Y. performed all the experiments with the help of T.Y.Y and L.X.F.; L.Y. and L.X.F. analyzed RNA-seq and mass spectrometry proteomics data; L.Q.C. and J.X.A. supervised the project; L.Y. and L.Q.C. wrote the manuscript. All authors contributed to data analysis and writing.

## Data and code availability

The RNA-seq data have been deposited in the NCBI Sequence Read Archive under accession number PRJNA836752. The mass spectrometry proteomics data have been deposited in the ProteomeXchange under accession number PXD033796.

## MATERIAS AND METHODS

### Bacterial strains and growth conditions

The bacterial strains, phage, and plasmids used in the present study are listed in Table S6. The *S. aureus* RN4220 or TZ0912 and *E. coli* IM08B were cultured in tryptic soy broth (TSB) and LB Broth media, respectively at 37°C. When required, the medium was supplemented with ampicillin (100 µg/mL) for *E. coli* and chloramphenicol (15 µg/mL) for *S. aureus* to maintain the pRAB11 or pOS1-*lacZ* plasmid. The pRAB11 plasmid was induced with 100 ng/mL anhydrotetracycline (ATc). The pIMAY plasmids used to construct mutants were selected using 25 µg/mL chloramphenicol for *E. coli* and 15 µg/mL chloramphenicol for *S. aureus*.

### Construction of mutants and plasmids

The oligonucleotides used in this study are listed in Table S5. The *S. aureus* TZ0912 mutant strains were generated by the allelic replacement method using temperature‐sensitive pIMAY plasmids as described previously ^64^. Briefly, ∼1000 bp DNA fragments flanking the genes of interest were amplified using the corresponding primer pairs USR_fw/ USR_rv and DSR_fw/ DSR_rv, and subsequently cloned into the pIMAY plasmid using NEBuilder HiFi DNA Assembly Master Mix. The resulting plasmids were transformed into *E. coli* IM08B and then electroporated into *S. aureus* TZ0912 followed by plating on TSA containing 15 µg/mL chloramphenicol at 28°C. Allelic exchange was performed as previously described ^64^. All mutants were verified by PCR amplification and sequencing. To construct the CRISPR-targeted plasmid, the protospacer sequence was annealed and cloned into the pRMC2 plasmid between the restriction sites Kpn I and Bgl II to obtain pCR1SP1. To construct plasmids containing the transcriptional fusions P*leader-lacZ and* P*cas-lacZ*, the P*leader* and putative P*cas* sequences were amplified by PCR from *S. aureus* TZ0912. The generated PCR products were cloned into the pOS1-*lacZ* plasmid digested with BamH I and EcoR I. The pET-AgrA, pET-SarA, pET- SarR and pET-ArcR plasmids used for purification of AgrA-His_6_, SarA-His_6_, SarR-His_6_ and ArcR-His_6_ proteins were generated by cloning the coding regions of the corresponding genes into the pET28a plasmid digested with Nde I and Xho I.

### Retention of Plasmid assay

*S. aureus* strains were transformed with the untargeted plasmid pRMC2 or CRISPR-targeted plasmid pCR1SP1. Single fresh *S. aureus* colonies of each transformation experiment with the pRMC2 or pCR1SP1 plasmid were picked and resuspended in 50 µL of TSB. Triplicates of 5 µL were inoculated in 5 mL of TSB with 100 ng/mL ATc and then incubated at 37ºC for 10 h. After incubation, the cultures were serially diluted and plated on TSB agar and TSB agar with 15 μg/mL chloramphenicol and incubated overnight at 37ºC before enumerating the CFUs. % CFUs containing plasmid was scored as the number of Cm^R^ CFU divided by the total number of CFU.

### Plaque assay

*S. aureus* overnight cultures (100 µL) were added to soft TSB agar (0.5% agar) and poured onto TSA plates. After solidification of the top agar, 5 µL of serially diluted phages was spotted onto the top agar and incubated overnight at 37°C. Full-plate assays were used to count plaques. Briefly, 25 µL of phage dilution (approximately 250 PFU) was incubated with 50 µL of *S. aureus* overnight culture for 10 min at 37 °C. Soft TSB agar (5 mL) was then added and the mixture was poured onto TSA agar. Individual plaques were counted and the efficiency of plaquing (EOP) was calculated.

### Growth curve

Overnight cultures of *S. aureus* were diluted to an OD_600_ of 0.05 in TSB with 5 mM CaCl_2_ and infected with phage phiIPLA-RODI at a multiplicity of infection (MOI) of 0.1 in biological triplicates in a 96- well plate. The plate was incubated at 37°C with shaking for 24 h and the OD_600_ was measured every 30 min using a microplate reader (TECAN, Spark).

### Expression and purification of proteins

AgrA-His_6_, SarA-His_6_, SarR-His_6_ and ArcR-His_6_ were expressed in *E. coli* BL21(DE3) cells. The cells were grown in 1 liter of LB broth at 37°C with shaking. At an optical density (OD_600_) of 0.4-0.6, 0.5 mM IPTG (isopropyl-β-D-thiogalactopyranoside) was added to induce protein expression. The cultures were incubated for an additional 16 h at 15°C with shaking, harvested by centrifugation, resuspended in phosphate buffered saline (137 mM NaCl, 2.7 mM KCl, 8.0 mM Na_2_HPO_4_, 1.5 mM KH_2_PO_4_, pH 7.4), and followed by sonication. After centrifugation, the clarified lysates were purified using an Ni^2+^-NTA-agarose affinity column (Takara) according to the manufacturer’s protocol.

### Electrophoretic mobility shift assays (EMSAs)

EMSAs were performed as follows. The labeled and unlabeled P*leader*, P*cas*, P*sarA*, and P*arcR* sequences were amplified with the primer pairs PE001/PE002, PE003/PE002, PE017/PE018, PE019/PE018, PE020/PE021, PE021/PE020, PE023/PE024 and PE025/PE024, respectively, using *S. aureus* TZ0912 chromosomal DNA as the template. The DNA fragment P2 promoter, which was used as a positive control, was annealed using the primer pair PE005/PE006. As a negative control probe, the house-keeping gene *gmk* intragenic DNA of *S. aureus* TZ0912 was also amplified with the primer pair PE007/PE008 by PCR. The PCR products were labeled using FAM-5’-end-labeled primers and then gel-purified using the MiniBEST Agarose Gel DNA Extraction kit (Takara). The binding reaction was carried out by incubating the labeled probe (20 ng) with increasing concentrations of the protein for 30 min at 37°C in a 20 μL solution containing the binding buffer (750mM NaCl, 50mM Tris-HCl (pH 7.4), 0.5mM DTT, 0.5mM EDTA). Protein-DNA binding reactions were resolved in a 6% non- denaturing acrylamide gel in 0.5 × TBE buffer, and images were scanned using a Typhoon FLA 9500 (GE Healthcare).

### β-galactosidase assays

The β-galactosidase activity was determined using o-nitrophenyl-β-D-galactopyranoside (ONPG) as described previously ^65^. Briefly, overnight cultures of *S. aureus* strains containing the P*leader-lacZ* or P*cas*-*lacZ* reporter were diluted to an OD_600_ of 0.05 in 5 mL TSB and then incubated at 37°C with shaking for 10 h. The OD_600_ of the cultures were measured, and 1 mL of the cultures were harvested by centrifugation. The cells were resuspended in 1 mL of Z buffer (60 mM Na_2_HPO_4_, 40 mM NaH_2_PHO_4_, 10 mM KCl, and 1 mM MgSO_4_, pH 7.0). The cells were incubated for an additional 15 min at 37°C with lysostaphin (1 mg/mL), after which 0.2 mL of ONPG (4 g/ml) was added and incubated at 37°C until the solution turned yellow. 0.4 mL of 1 M Na_2_CO_3_ was added to stop the enzymatic reactions, and the OD_420_ was measured. The β-galactosidase activity was calculated as Miller Units. The assay was repeated at least three times.

### Total RNA extraction and qRT-PCR

Overnight cultures of *S. aureus* strains were diluted to an OD_600_ of 0.05 in 30 mL TSB and then incubated at 37°C with shaking. Cells were harvested by centrifugation at the indicated time points. The cells were lysed with lysostaphin, and total RNA was extracted using the RNeasy Mini Kit according to the manufacturer’s instructions (Qiagen). The removal of genomic DNA (gDNA) and reverse transcription reactions were performed using the PrimeScript RT Regent Kit with gDNA Eraser (Takara). cDNA was quantified by qRT-PCR using FastStart Universal SYBR Green Master (ROX) (Roche). The *gyrB* gene was used as an endogenous control for normalization. Each qRT-PCR analysis was performed with at least three biological replicates.

### RNA-Seq

RNA libraries were prepared with a TruSeq RNA Sample Prep Kit (Illumina) using 3 μg of total RNA extracted from cultures grown for 9 h and then sequenced on an Illumina HiSeq4000 sequencer. RNA- seq reads were aligned to the reference genome *S. aureus* TZ0912 using Bowtie2.

### RT-PCR

A total of 2 μL of the cDNA product was used as a template for subsequent PCR using primer pairs that amplify regions spanning different combinations of genes. As negative and positive controls, the RNA with gDNA removed and gDNA were used as templates in the PCR reaction, respectively.

### DNA pull-down

DNA pull-down was performed as described previously, with modifications to the protocol ^66^. In brief, the P*leader* and P*cas* sequences were amplified with the 5’-biotin-labeled primer pairs PD001/PE002 and PD005/PE018 by PCR using *S. aureus* TZ0912 chromosomal DNA as the template, respectively. As a negative control probe, the house-keeping gene *gmk* intragenic DNA of *S. aureus* TZ0912 was also amplified with the 5’-biotin-labeled primer pair PD003/PE008 by PCR. The resulting PCR products were gel-purified using the MiniBEST Agarose Gel DNA Extraction kit (Takara). Overnight cultures of *S. aureus* TZ0912 were diluted to an OD_600_ of 0.05 in 1 L TSB and then incubated at 37°C with shaking for 10 h. The cells were harvested by centrifugation, resuspended in BS/THES buffer (22 mM Tris-HCl pH=7.5, 4.4 mM EDTA, 8.9% (w/v) sucrose, 62 mM NaCl, 0.7% Protease Inhibitor Cocktail (v/v), 10 mM HEPES, 5 mM CaCl_2_, 50 mM KCl, 12% glycerol). After treatment with lysostaphin at 37°C for 1 h, the sample was lysed by sonication. The clarified lysate was obtained by centrifuging twice. The 200 μL Dynabeads M-280 Streptavidin (Invitrogen) was washed twice with 500 μL 2×B/W buffer (10 mM Tris HCl pH 7.5, 1 mM EDTA, 2 M NaCl) before incubating the beads and 40 μg DNA probe at 25°C with shaking for 1 h. The bead-DNA complex was washed twice with 400 μL TE buffer (0.5 M Tris-HCl, pH 8.0; 1 mM EDTA, pH 8.0) and then twice with 500 μL BS/THES buffer. The bead-DNA was resuspended in 500 μL of BS/THES buffer supplemented with 10 μg/mL poly(dI-dC) (Sigma) and mixed with 700 μL clarified lysate at 25°C for 30 min. The bead-DNA was washed five times with BS/THES buffer containing 10 μg/mL poly(dI-dC), and then twice with BS/THES. The bound proteins were eluted with increasing NaCl concentrations (0.1, 0.2, 0.3, 0.5, 0.75, 1 M). The eluates were subjected to SDS-PAGE followed by silver staining and mass spectrometry.

### Phylogenetic analysis

To identify the P*leader* and P*cas* homologs in Firmicutes, the sequences were used as queries against the NCBI nr nucleotide database. Hits with sequence identity >65% and query coverage >35% were retained for further analysis. The representative sequences were aligned using Clustal Omega, and phylogenetic trees were constructed using the neighbor-joining clustering method. Branch support was computed using 1,000 bootstrap replicates. The tree data were visualized and modified using iTOL software.

## REFERENCES

1. Hampton, H.G., Watson, B.N.J., and Fineran, P.C. (2020). The arms race between bacteria and their phage foes. Nature 577, 327–336. 10.1038/s41586-019-1894-8.

2. Barrangou, R., Fremaux, C., Deveau, H., Richards, M., Boyaval, P., Moineau, S., Romero, D.A., and Horvath, P. (2007). CRISPR Provides Acquired Resistance Against Viruses in Prokaryotes. Science 315, 1709–1712. 10.1126/science.1138140.

3. Marraffini, L.A., and Sontheimer, E.J. (2008). CRISPR Interference Limits Horizontal Gene Transfer in Staphylococci by Targeting DNA. Science 322, 1843–1845. 10.1126/science.1165771.

4. Makarova, K.S., Wolf, Y.I., Iranzo, J., Shmakov, S.A., Alkhnbashi, O.S., Brouns, S.J.J., Charpentier, E., Cheng, D., Haft, D.H., Horvath, P., et al. (2020). Evolutionary classification of CRISPR–Cas systems: a burst of class 2 and derived variants. Nature Reviews Microbiology 18, 67–83. 10.1038/s41579-019-0299-x.

5. Marraffini, L.A. (2015). CRISPR-Cas immunity in prokaryotes. Nature 526, 55–61. 10.1038/nature15386.

6. Nuñez, J.K., Lee, A.S.Y., Engelman, A., and Doudna, J.A. (2015). Integrase-mediated spacer acquisition during CRISPR–Cas adaptive immunity. Nature 519, 193–198. 10.1038/nature14237.

7. Yosef, I., Goren, M.G., and Qimron, U. (2012). Proteins and DNA elements essential for the CRISPR adaptation process in Escherichia coli. Nucleic Acids Research 40, 5569–5576. 10.1093/nar/gks216.

8. Carte, J., Wang, R., Li, H., Terns, R.M., and Terns, M.P. (2008). Cas6 is an endoribonuclease that generates guide RNAs for invader defense in prokaryotes. Genes Dev 22, 3489–3496. 10.1101/gad.1742908.

9. Brouns, S.J., Jore, M.M., Lundgren, M., Westra, E.R., Slijkhuis, R.J., Snijders, A.P., Dickman, M.J., Makarova, K.S., Koonin, E.V., and van der Oost, J. (2008). Small CRISPR RNAs guide antiviral defense in prokaryotes. Science 321, 960–964. 10.1126/science.1159689.

10. Garneau, J.E., Dupuis, M.E., Villion, M., Romero, D.A., Barrangou, R., Boyaval, P., Fremaux, C., Horvath, P., Magadan, A.H., and Moineau, S. (2010). The CRISPR/Cas bacterial immune system cleaves bacteriophage and plasmid DNA. Nature 468, 67–71. 10.1038/nature09523.

11. Goldberg, G.W., Jiang, W., Bikard, D., and Marraffini, L.A. (2014). Conditional tolerance of temperate phages via transcription-dependent CRISPR-Cas targeting. Nature 514, 633–637. 10.1038/nature13637.

12. Samai, P., Pyenson, N., Jiang, W., Goldberg, Gregory W., Hatoum-Aslan, A., and Marraffini, Luciano A. (2015). Co-transcriptional DNA and RNA Cleavage during Type III CRISPR-Cas Immunity. Cell 161, 1164–1174. 10.1016/j.cell.2015.04.027.

13. Deng, L., Garrett, R.A., Shah, S.A., Peng, X., and She, Q. (2013). A novel interference mechanism by a type IIIB CRISPR-Cmr module in Sulfolobus. Mol Microbiol 87, 1088–1099. 10.1111/mmi.12152.

14. Elmore, J.R., Sheppard, N.F., Ramia, N., Deighan, T., Li, H., Terns, R.M., and Terns, M.P. (2016). Bipartite recognition of target RNAs activates DNA cleavage by the Type III-B CRISPR-Cas system. Genes Dev 30, 447–459. 10.1101/gad.272153.115.

15. Estrella, M.A., Kuo, F.T., and Bailey, S. (2016). RNA-activated DNA cleavage by the Type III-B CRISPR-Cas effector complex. Genes Dev 30, 460–470. 10.1101/gad.273722.115.

16. Kazlauskiene, M., Tamulaitis, G., Kostiuk, G., Venclovas, C., and Siksnys, V. (2016). Spatiotemporal Control of Type III-A CRISPR-Cas Immunity: Coupling DNA Degradation with the Target RNA Recognition. Mol Cell 62, 295–306. 10.1016/j.molcel.2016.03.024.

17. Kazlauskiene, M., Kostiuk, G., Venclovas, Č., Tamulaitis, G., and Siksnys, V. (2017). A cyclic oligonucleotide signaling pathway in type III CRISPR-Cas systems. Science 357, 605–609. 10.1126/science.aao0100.

18. Niewoehner, O., Garcia-Doval, C., Rostol, J.T., Berk, C., Schwede, F., Bigler, L., Hall, J., Marraffini, L.A., and Jinek, M. (2017). Type III CRISPR-Cas systems produce cyclic oligoadenylate second messengers. Nature 548, 543–548. 10.1038/nature23467.

19. Li, Q., Xie, X., Yin, K., Tang, Y., Zhou, X., Chen, Y., Xia, J., Hu, Y., Ingmer, H., Li, Y., and Jiao, X. (2016). Characterization of CRISPR-Cas system in clinical Staphylococcus epidermidis strains revealed its potential association with bacterial infection sites. Microbiological Research 193, 103–110. 10.1016/j.micres.2016.09.003.

20. Li, Y., Mikkelsen, K., Lluch, I.G.O., Wang, Z., Tang, Y., Jiao, X., Ingmer, H., Hoyland-Kroghsbo, N.M., and Li, Q. (2021). Functional Characterization of Type III-A CRISPR-Cas in a Clinical Human Methicillin-R Staphylococcus aureus Strain. CRISPR J 4, 686–698. 10.1089/crispr.2021.0046.

21. Cao, L., Gao, C.-H., Zhu, J., Zhao, L., Wu, Q., Li, M., and Sun, B. (2016). Identification and functional study of type III-A CRISPR-Cas systems in clinical isolates of Staphylococcus aureus. International Journal of Medical Microbiology 306, 686–696. 10.1016/j.ijmm.2016.08.005.

22. Patterson, A.G., Jackson, S.A., Taylor, C., Evans, G.B., Salmond, G.P.C., Przybilski, R., Staals, R.H.J., and Fineran, P.C. (2016). Quorum Sensing Controls Adaptive Immunity through the Regulation of Multiple CRISPR-Cas Systems. Molecular Cell 64, 1102–1108. 10.1016/j.molcel.2016.11.012.

23. Høyland-Kroghsbo, N.M., Paczkowski, J., Mukherjee, S., Broniewski, J., Westra, E., Bondy-Denomy, J., and Bassler, B.L. (2017). Quorum sensing controls the Pseudomonas aeruginosa CRISPR-Cas adaptive immune system. Proceedings of the National Academy of Sciences 114, 131–135. 10.1073/pnas.1617415113.

24. Mukherjee, S., and Bassler, B.L. (2019). Bacterial quorum sensing in complex and dynamically changing environments. Nature Reviews Microbiology 17, 371–382. 10.1038/s41579-019-0186-5.

25. Gao, R., Krysciak, D., Petersen, K., Utpatel, C., Knapp, A., Schmeisser, C., Daniel, R., Voget, S., Jaeger, K.E., and Streit, W.R. (2015). Genome-wide RNA sequencing analysis of quorum sensing-controlled regulons in the plant-associated Burkholderia glumae PG1 strain. Appl Environ Microbiol 81, 7993–8007. 10.1128/AEM.01043-15.

26. Kasman, L.M., Kasman, A., Westwater, C., Dolan, J., Schmidt, M.G., and Norris, J.S. (2002). Overcoming the phage replication threshold: a mathematical model with implications for phage therapy. J Virol 76, 5557–5564. 10.1128/jvi.76.11.5557-5564.2002.

27. Painter, K.L., Krishna, A., Wigneshweraraj, S., and Edwards, A.M. (2014). What role does the quorum-sensing accessory gene regulator system play during Staphylococcus aureus bacteremia? Trends Microbiol 22, 676–685. 10.1016/j.tim.2014.09.002.

28. Bronesky, D., Wu, Z., Marzi, S., Walter, P., Geissmann, T., Moreau, K., Vandenesch, F., Caldelari, I., and Romby, P. (2016). Staphylococcus aureus RNAIII and Its Regulon Link Quorum Sensing, Stress Responses, Metabolic Adaptation, and Regulation of Virulence Gene Expression. Annu Rev Microbiol 70, 299–316. 10.1146/annurev-micro-102215-095708.

29. Queck, S.Y., Jameson-Lee, M., Villaruz, A.E., Bach, T.H., Khan, B.A., Sturdevant, D.E., Ricklefs, S.M., Li, M., and Otto, M. (2008). RNAIII-independent target gene control by the agr quorum-sensing system: insight into the evolution of virulence regulation in Staphylococcus aureus. Mol Cell 32, 150–158. 10.1016/j.molcel.2008.08.005.

30. Makhlin, J., Kofman, T., Borovok, I., Kohler, C., Engelmann, S., Cohen, G., and Aharonowitz, Y. (2007). Staphylococcus aureus ArcR controls expression of the arginine deiminase operon. J Bacteriol 189, 5976–5986. 10.1128/JB.00592-07.

31. Cameron, A.D., and Redfield, R.J. (2006). Non-canonical CRP sites control competence regulons in Escherichia coli and many other gamma-proteobacteria. Nucleic Acids Res 34, 6001–6014. 10.1093/nar/gkl734.

32. Sterba, K.M., Mackintosh, S.G., Blevins, J.S., Hurlburt, B.K., and Smeltzer, M.S. (2003). Characterization of Staphylococcus aureus SarA binding sites. J Bacteriol 185, 4410–4417. 10.1128/JB.185.15.4410-4417.2003.

33. Oriol, C., Cengher, L., Manna, A.C., Mauro, T., Pinel-Marie, M.L., Felden, B., Cheung, A., and Rouillon, A. (2021). Expanding the Staphylococcus aureus SarA Regulon to Small RNAs. mSystems 6, e0071321. 10.1128/mSystems.00713-21.

34. Mauro, T., Rouillon, A., and Felden, B. (2016). Insights into the regulation of small RNA expression: SarA represses the expression of two sRNAs in Staphylococcus aureus. Nucleic Acids Res 44, 10186–10200. 10.1093/nar/gkw777.

35. Reyes, D., Andrey, D.O., Monod, A., Kelley, W.L., Zhang, G., and Cheung, A.L. (2011). Coordinated regulation by AgrA, SarA, and SarR to control agr expression in Staphylococcus aureus. J Bacteriol 193, 6020–6031. 10.1128/JB.05436-11.

36. Dunman, P.M., Murphy, E., Haney, S., Palacios, D., Tucker-Kellogg, G., Wu, S., Brown, E.L., Zagursky, R.J., Shlaes, D., and Projan, S.J. (2001). Transcription profiling-based identification of Staphylococcus aureus genes regulated by the agr and/or sarA loci. J Bacteriol 183, 7341–7353. 10.1128/JB.183.24.7341-7353.2001.

37. Koenig, R.L., Ray, J.L., Maleki, S.J., Smeltzer, M.S., and Hurlburt, B.K. (2004). Staphylococcus aureus AgrA binding to the RNAIII-agr regulatory region. J Bacteriol 186, 7549–7555. 10.1128/JB.186.22.7549-7555.2004.

38. Hoyland-Kroghsbo, N.M., Maerkedahl, R.B., and Svenningsen, S.L. (2013). A quorum-sensing-induced bacteriophage defense mechanism. mBio 4, e00362–00312. 10.1128/mBio.00362-12.

39. McGinn, J., and Marraffini, Luciano A. (2016). CRISPR-Cas Systems Optimize Their Immune Response by Specifying the Site of Spacer Integration. Molecular Cell 64, 616–623. 10.1016/j.molcel.2016.08.038.

40. Ali, L., Goraya, M.U., Arafat, Y., Ajmal, M., Chen, J.L., and Yu, D. (2017). Molecular Mechanism of Quorum-Sensing in Enterococcus faecalis: Its Role in Virulence and Therapeutic Approaches. Int J Mol Sci 18. 10.3390/ijms18050960.

41. Suntharalingam, P., and Cvitkovitch, D.G. (2005). Quorum sensing in streptococcal biofilm formation. Trends Microbiol 13, 3–6. 10.1016/j.tim.2004.11.009.

42. Przybilski, R., Richter, C., Gristwood, T., Clulow, J.S., Vercoe, R.B., and Fineran, P.C. (2011). Csy4 is responsible for CRISPR RNA processing in Pectobacterium atrosepticum. RNA Biol 8, 517–528. 10.4161/rna.8.3.15190.

43. Pul, U., Wurm, R., Arslan, Z., Geissen, R., Hofmann, N., and Wagner, R. (2010). Identification and characterization of E. coli CRISPR-cas promoters and their silencing by H-NS. Mol Microbiol 75, 1495–1512. 10.1111/j.1365-2958.2010.07073.x.

44. Wei, Y., Chesne, M.T., Terns, R.M., and Terns, M.P. (2015). Sequences spanning the leader-repeat junction mediate CRISPR adaptation to phage in Streptococcus thermophilus. Nucleic Acids Res 43, 1749–1758. 10.1093/nar/gku1407.

45. Hatoum-Aslan, A., Maniv, I., Samai, P., and Marraffini, L.A. (2014). Genetic Characterization of Antiplasmid Immunity through a Type III-A CRISPR-Cas System. Journal of Bacteriology 196, 310–317. 10.1128/jb.01130-13.

46. Patterson, A.G., Yevstigneyeva, M.S., and Fineran, P.C. (2017). Regulation of CRISPR-Cas adaptive immune systems. Curr Opin Microbiol 37, 1–7. 10.1016/j.mib.2017.02.004.

47. Bernal-Bernal, D., Abellon-Ruiz, J., Iniesta, A.A., Pajares-Martinez, E., Bastida-Martinez, E., Fontes, M., Padmanabhan, S., and Elias-Arnanz, M. (2018). Multifactorial control of the expression of a CRISPR-Cas system by an extracytoplasmic function sigma/anti-sigma pair and a global regulatory complex. Nucleic Acids Res 46, 6726–6745. 10.1093/nar/gky475.

48. Chien, Y., Manna, A.C., Projan, S.J., and Cheung, A.L. (1999). SarA, a global regulator of virulence determinants in Staphylococcus aureus, binds to a conserved motif essential for sar-dependent gene regulation. J Biol Chem 274, 37169–37176. 10.1074/jbc.274.52.37169.

49. Schilcher, K., and Horswill, A.R. (2020). Staphylococcal Biofilm Development: Structure, Regulation, and Treatment Strategies. Microbiol Mol Biol Rev 84, e00026–00019. 10.1128/MMBR.00026-19.

50. Schuch, R., Khan, B.K., Raz, A., Rotolo, J.A., and Wittekind, M. (2017). Bacteriophage Lysin CF-301, a Potent Antistaphylococcal Biofilm Agent. Antimicrob Agents Chemother 61. 10.1128/AAC.02666-16.

51. Alves, D.R., Gaudion, A., Bean, J.E., Perez Esteban, P., Arnot, T.C., Harper, D.R., Kot, W., Hansen, L.H., Enright, M.C., and Jenkins, A.T. (2014). Combined use of bacteriophage K and a novel bacteriophage to reduce Staphylococcus aureus biofilm formation. Appl Environ Microbiol 80, 6694–6703. 10.1128/AEM.01789-14.

52. Manna, A., and Cheung, A.L. (2001). Characterization of sarR, a modulator of sar expression in Staphylococcus aureus. Infect Immun 69, 885–896. 10.1128/IAI.69.2.885-896.2001.

53. You, C., Okano, H., Hui, S., Zhang, Z., Kim, M., Gunderson, C.W., Wang, Y.P., Lenz, P., Yan, D., and Hwa, T. (2013). Coordination of bacterial proteome with metabolism by cyclic AMP signalling. Nature 500, 301–306. 10.1038/nature12446.

54. Yang, C.D., Chen, Y.H., Huang, H.Y., Huang, H.D., and Tseng, C.P. (2014). CRP represses the CRISPR/Cas system in Escherichia coli: evidence that endogenous CRISPR spacers impede phage P1 replication. Mol Microbiol 92, 1072–1091. 10.1111/mmi.12614.

55. Patterson, A.G., Chang, J.T., Taylor, C., and Fineran, P.C. (2015). Regulation of the Type I-F CRISPR-Cas system by CRP-cAMP and GalM controls spacer acquisition and interference. Nucleic Acids Research 43, 6038–6048. 10.1093/nar/gkv517.

56. Shinkai, A., Kira, S., Nakagawa, N., Kashihara, A., Kuramitsu, S., and Yokoyama, S. (2007). Transcription activation mediated by a cyclic AMP receptor protein from Thermus thermophilus HB8. J Bacteriol 189, 3891–3901. 10.1128/JB.01739-06.

57. Agari, Y., Sakamoto, K., Tamakoshi, M., Oshima, T., Kuramitsu, S., and Shinkai, A. (2010). Transcription profile of Thermus thermophilus CRISPR systems after phage infection. J Mol Biol 395, 270–281. 10.1016/j.jmb.2009.10.057.

58. Tan, D., Svenningsen, S.L., and Middelboe, M. (2015). Quorum Sensing Determines the Choice of Antiphage Defense Strategy in Vibrio anguillarum. mBio 6, e00627. 10.1128/mBio.00627-15.

59. Depardieu, F., Didier, J.P., Bernheim, A., Sherlock, A., Molina, H., Duclos, B., and Bikard, D. (2016). A Eukaryotic-like Serine/Threonine Kinase Protects Staphylococci against Phages. Cell Host Microbe 20, 471–481. 10.1016/j.chom.2016.08.010.

60. Smith, L.M., Jackson, S.A., Malone, L.M., Ussher, J.E., Gardner, P.P., and Fineran, P.C. (2021). The Rcs stress response inversely controls surface and CRISPR-Cas adaptive immunity to discriminate plasmids and phages. Nat Microbiol 6, 162–172. 10.1038/s41564-020-00822-7.

61. Workman, R.E., Pammi, T., Nguyen, B.T.K., Graeff, L.W., Smith, E., Sebald, S.M., Stoltzfus, M.J., Euler, C.W., and Modell, J.W. (2021). A natural single-guide RNA repurposes Cas9 to autoregulate CRISPR-Cas expression. Cell 184, 675–688 e619. 10.1016/j.cell.2020.12.017.

62. Campa, A.R., Smith, L.M., Hampton, H.G., Sharma, S., Jackson, S.A., Bischler, T., Sharma, C.M., and Fineran, P.C. (2021). The Rsm (Csr) post-transcriptional regulatory pathway coordinately controls multiple CRISPR-Cas immune systems. Nucleic Acids Res. 10.1093/nar/gkab704.

63. Borges, A.L., Castro, B., Govindarajan, S., Solvik, T., Escalante, V., and Bondy-Denomy, J. (2020). Bacterial alginate regulators and phage homologs repress CRISPR–Cas immunity. Nature Microbiology 5, 679–687. 10.1038/s41564-020-0691-3.

64. Monk, I.R., Shah, I.M., Xu, M., Tan, M.-W., Foster, T.J., and Novick, R.P. (2012). Transforming the Untransformable: Application of Direct Transformation To Manipulate Genetically Staphylococcus aureus and Staphylococcus epidermidis. mBio 3. 10.1128/mBio.00277-11.

65. Smale, S.T. (2010). Beta-galactosidase assay. Cold Spring Harb Protoc 2010, pdb prot5423. 10.1101/pdb.prot5423.

66. Chaparian, R.R., and van Kessel, J.C. (2021). Promoter Pull-Down Assay A Biochemical Screen for DNA-Binding Proteins. Methods in Molecular Biology 2346, 165–172. 10.1007/7651_2020_307.

