## Supplemental Figure S1-5 for "Quorum sensing inhibits Type III-A CRISPR-Cas system activity through repressing positive regulators SarA and ArcR in *Staphylococcus aureus*"

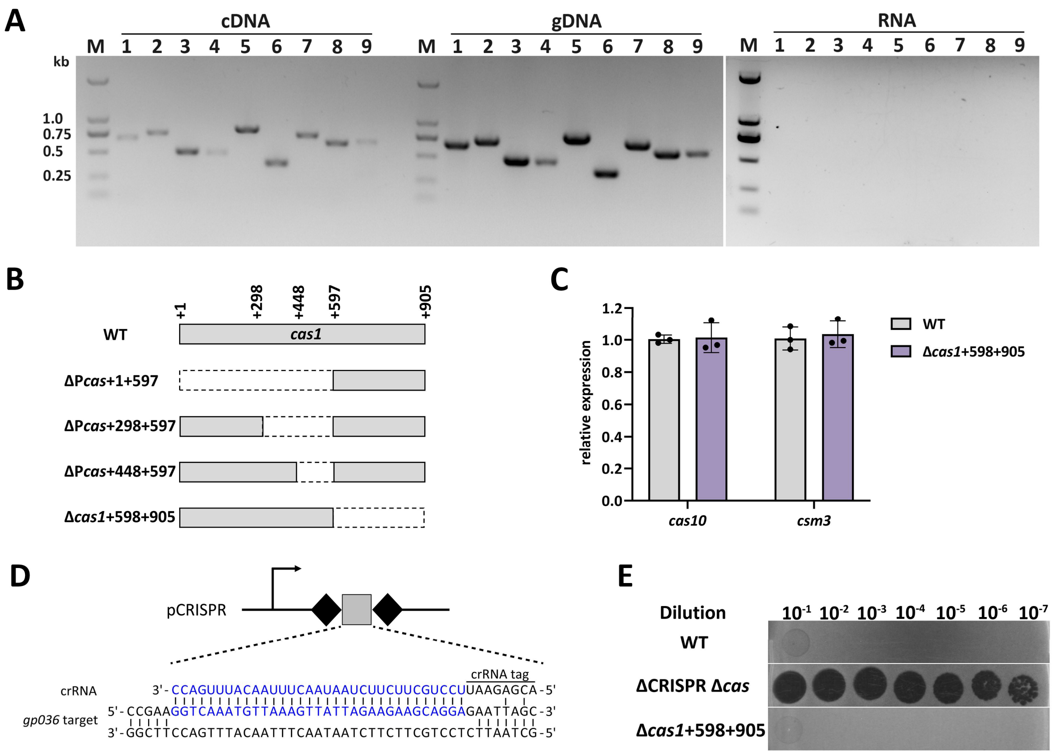


**Figure S1. P*cas* sequence drives the transcription of *cas* gene clusters.** (**A**) RT-PCR amplification of intergenic regions was conducted using cDNA, gDNA and RNA as templates, respectively. The amplified intergenic regions were indicated by numbers that correspond to the as black horizontal lines numbered 1–9 in Figure 1A. M, 2 kb DNA ladder. (**B**) Schematic presentation of the respective P*cas* mutants used in this study. The deleted region was indicted by dashed box. (**C**) qRT-PCR measurement of *cas10* and *csm3* expression in the WT and Δ*cas1* (308 bp) mutants after 10 h of growth in liquid culture, respectively. Data shown are means ± standard deviation of three independent experiments. (**D**) The pCRISPR plasmid encodes a spacer that targets the *gp036* gene of phage phiIPLA-RODI. The crRNA and target sequences are shown in blue. (**E**) Tenfold serial dilution of phage phiIPLA-RODI was spotted on the bacterial lawns of WT, ΔCRISPR Δ*cas* and Δ*cas1* (308 bp) mutant carrying the pCRISPR plasmid. Plaquing experiments were replicated three times and consistent results were seen.

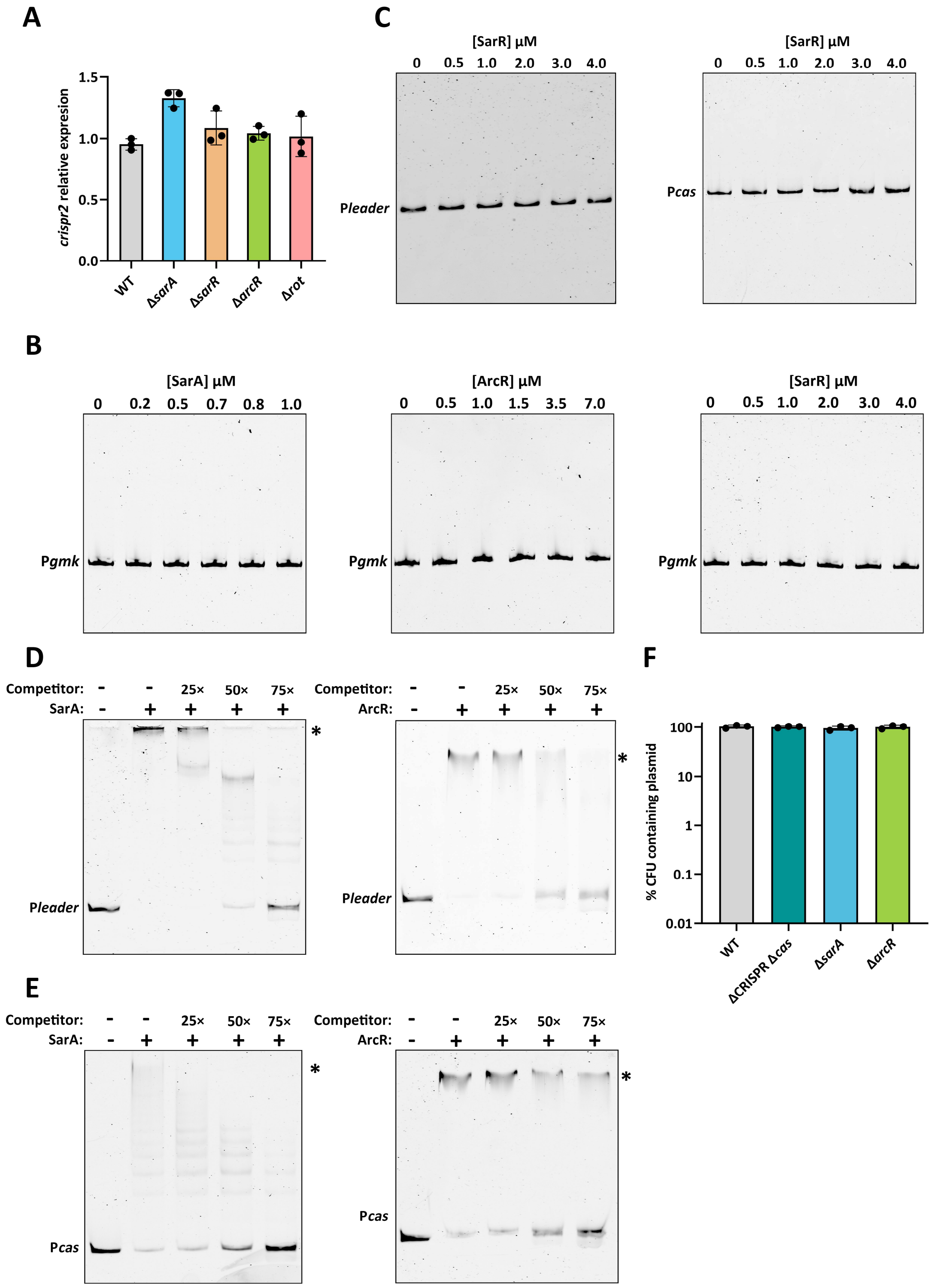


**Figure S2. SarA and ArcR cannot interact with negative probe.** (**A**) qRT-PCR analysis of *crispr2* expression in the WT and the individual mutant strains at high cell density, respectively. (**B**) FAM-5'-end-labelled DNA fragment of P*gmk* was incubated with increasing concentration of purified SarA-His$_6$, ArcR-His$_6$ and SarR-His$_6$ protein. The DNA fragment of *gmk* was used as a negative control. (**C**) FAM-5'-end-labelled P*leader* and P*cas* sequence was incubated with increasing concentration of purified SarR-His$_6$ protein, respectively. (**D**) Competitive EMSAs. FAM-5'-end-labelled P*leader* sequence was incubated with 1.0 μM purified SarA-His$_6$ and 3.0 μM purified ArcR-His$_6$ in the absence or presence of 25-, 50-, and 75- fold excess of unlabeled

P*leader* sequence competitors. DNA-protein complexes are indicated by an asterisk. (**E**) Same assay as in (D). FAM-5'-end-labelled P*cas* sequence was incubated with 1.0 μM purified SarA-$His_6$ and 7.0 μM purified ArcR-$His_6$ in the absence or presence of 25-, 50-, and 75- fold excess of unlabeled P*cas* sequence competitors. DNA-protein complexes are indicated by an asterisk.

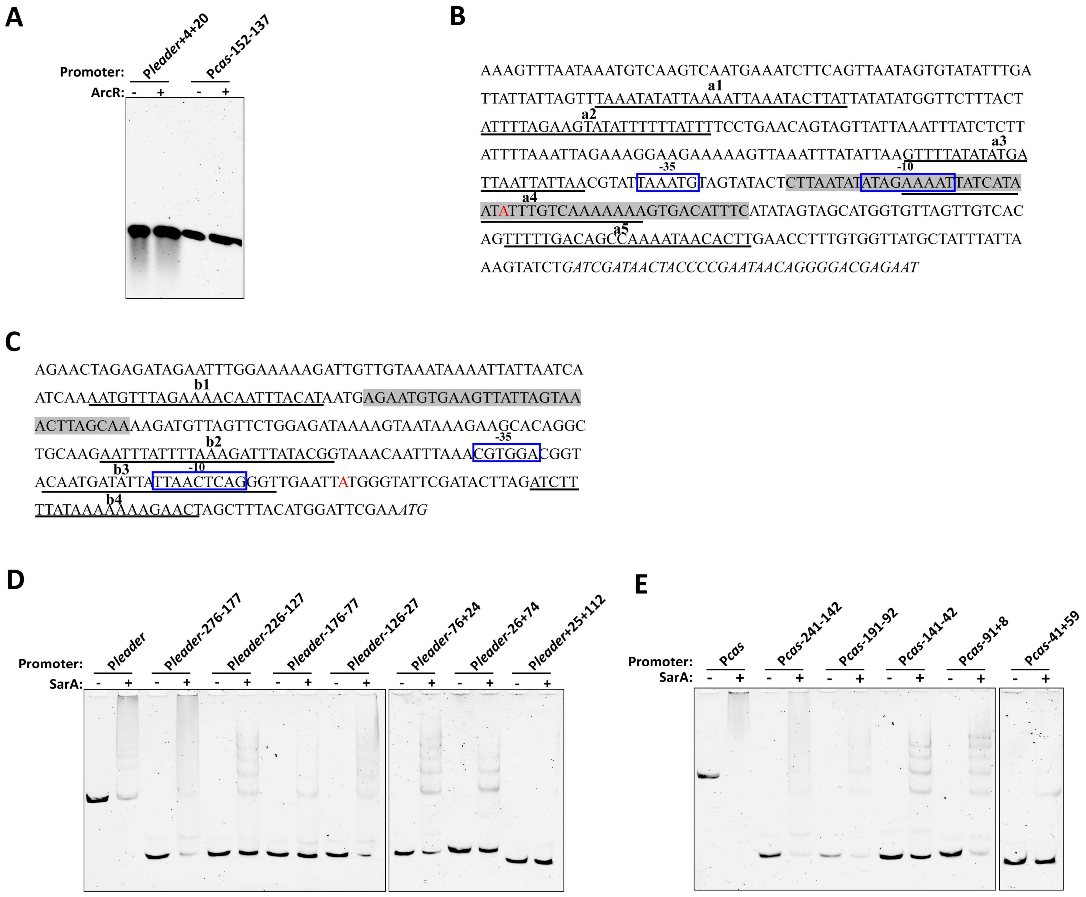


**Figure S3. Determination of ArcR and SarA binding sites in P*leader* and P*cas* sequence by EMSA.** (**A**) FAM-5'-end-labelled P*leader*+4+20 and P*cas*-152-137 fragment was incubated with ArcR-$His_6$ at the concentration of 7.2 µM, respectively. (**B**) Nucleotide sequence of the CRISPR1 leader sequence. Transcriptional start site predicted by RNA-seq is indicated in red. -10 and -35 regions, labeled in blue, are also boxed. ArcR binding site is shadowed in gray. SarA binding sites (a1-a5) are underlined. The first repeat of the CRISPR 1 array is marked in italic font. (**C**) Nucleotide sequence of the P*cas* sequence. Transcriptional start site predicted by RNA-seq is indicated in red. -10 and -35 regions, labeled in blue, are also boxed. ArcR binding site is shadowed in gray. SarA binding sites (b1-b4) are underlined. The ATG position from +598 to +600 of *cas1* is marked in italic font. (**D**) Seven truncated FAM-5'-end-labelled P*leader* fragments were incubated with SarA-$His_6$ at the concentration of 1.0 µM, respectively. (**E**) Five truncated FAM-5'-end-labelled P*cas* fragments were incubated with SarA-$His_6$ at the concentration of 1.0 µM, respectively. DNA-protein complexes are indicated by an asterisk. +, with protein; –, without protein. (**F**) RNA-seq reads were mapped to the WT and Δ*sarA* genome to determine the relative abundance of P*cas* sequence. Position of P*cas* sequence is indicated with vertical dotted lines.

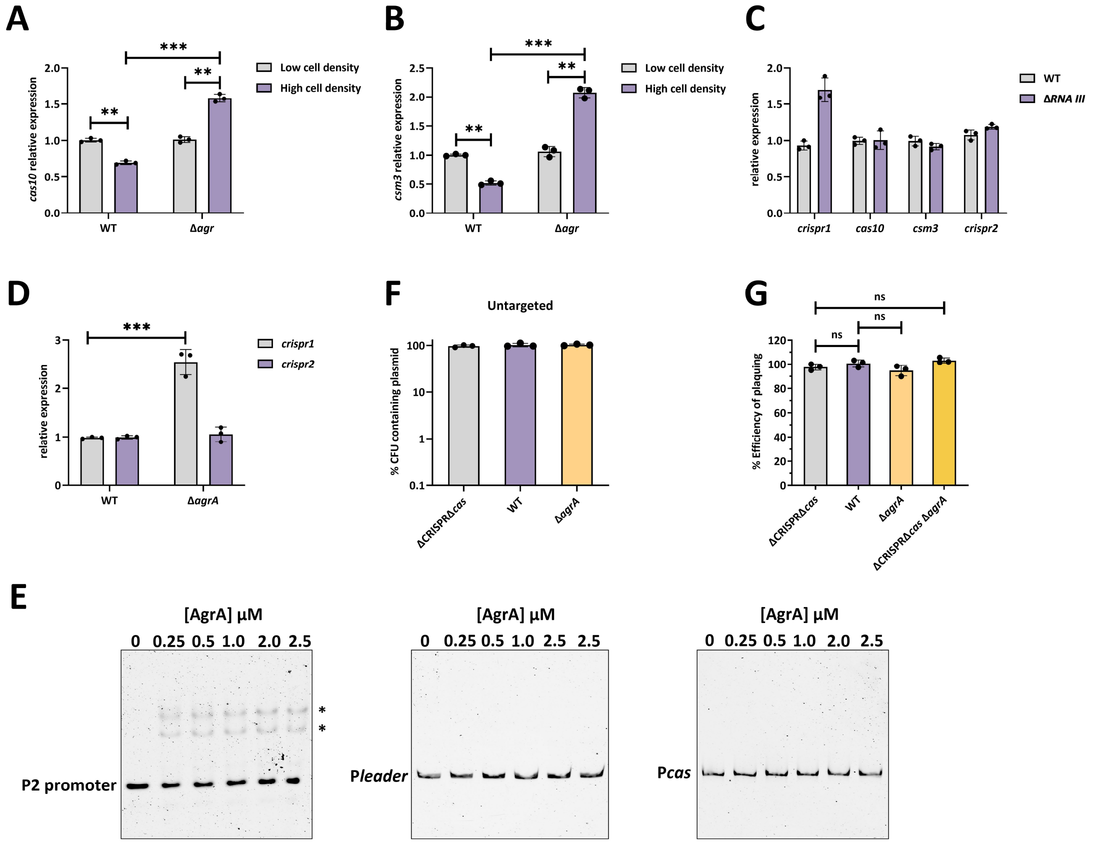


**Figure S4. The immunity against phage infection of QS mutant.** (**A**) The expression of *cas10* was measured by qRT-PCR in the WT and Δ*agr* mutant at both low and high cell densities, respectively. The *gyrB* gene was used as the internal control. (**B**) Same as (**A**), but the *csm3* expression was assessed. (**C**) qRT-PCR analysis of *crispr1*, *crispr2*, *cas10* and *csm3* expression in the WT and Δ*RNA III* mutant at high cell density. (**D**) qRT-PCR analysis of *crispr1* and *crispr2* expression in the WT and Δ*agrA* mutant at high cell density. (**E**) FAM-5'-end-labelled P*leader* and P*cas* DNA fragments were incubated with increasing concentrations of purified AgrA-His$_6$ protein. Acetyl phosphate (50 mM) was added to all EMSAs. The P2 promoter, a promoter of the *agr* QS system that is regulated directly by AgrA, was used as a positive control. DNA-protein complexes are indicated by an asterisk. (**F**) Retention of the untargeted plasmid pRMC2 in the WT, ΔCRISPR Δ*cas* and Δ*agrA* mutant. The strains containing the plasmid were grown in TSB with ATc for 10 h followed by plating on TSA with or without chloramphenicol. % CFUs containing plasmid was scored as the number of Cm$^R$ CFU/the total number of CFU. (**G**) The immunity against phage phiSA012 of relevant strains was represented as efficiency of plaquing, which is a ratio relative to the number of plaque measured on the ΔCRISPR Δ*cas* mutant. Data shown are means ± standard deviation of three independent experiments. A two-tailed unpaired Student's *t*-test was used to calculate *P* values; **$p$<0.01, ***$p$<0.001. ns, not significant.

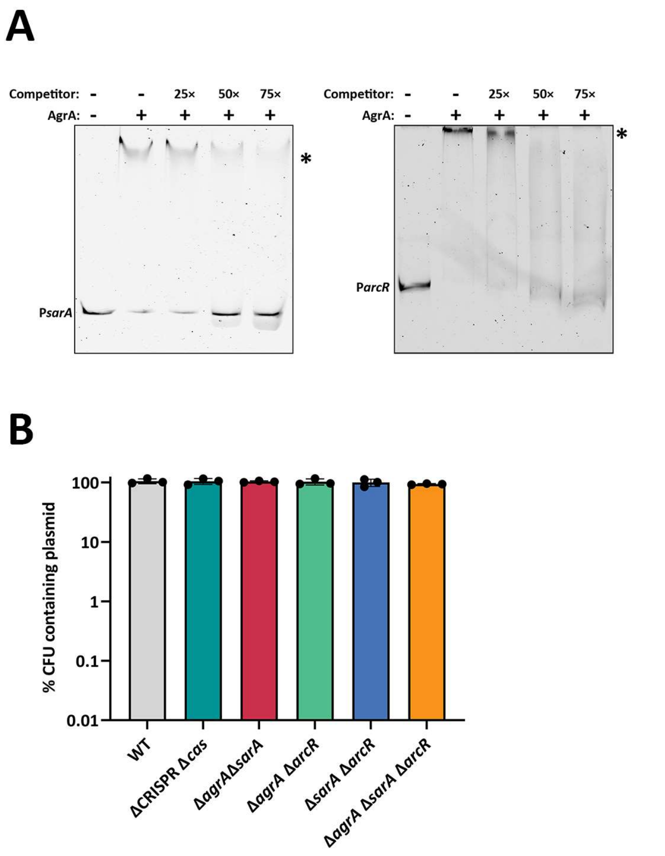


**Figure S5. AgrA specifically binds to the P*sarA* and P*arcR* sequence.** (**A**) Competitive EMSAs. FAM-5'-end-labelled P*sarA* sequence or P*arcR* sequence was incubated with 2 µM purified AgrA-$His_6$ protein in the absence or presence of 25-, 50-, and 75- fold excess of unlabeled P*sarA* sequence or P*arcR* sequence competitors, respectively. DNA-protein complexes are indicated by an asterisk. (**B**) Retention of the untargeted plasmid pRMC2 in the WT, ΔCRISPR Δ*cas*, and double mutants (Δ*agrA* Δ*sarA*, Δ*agrA* Δ*arcR,* Δ*sarA* Δ*arcR*), and triple mutant Δ*agrA* Δ*sarA* Δ*arcR*. The strains containing the pRMC2 plasmid were grown in TSB with ATc for 10 h followed by plating on TSA with or without chloramphenicol. % CFUs represents the ratio of bacteria grown on plates containing chloramphenicol to the number of bacteria growing on plates without antibiotics. Data shown are means ± standard deviation of three independent experiment.

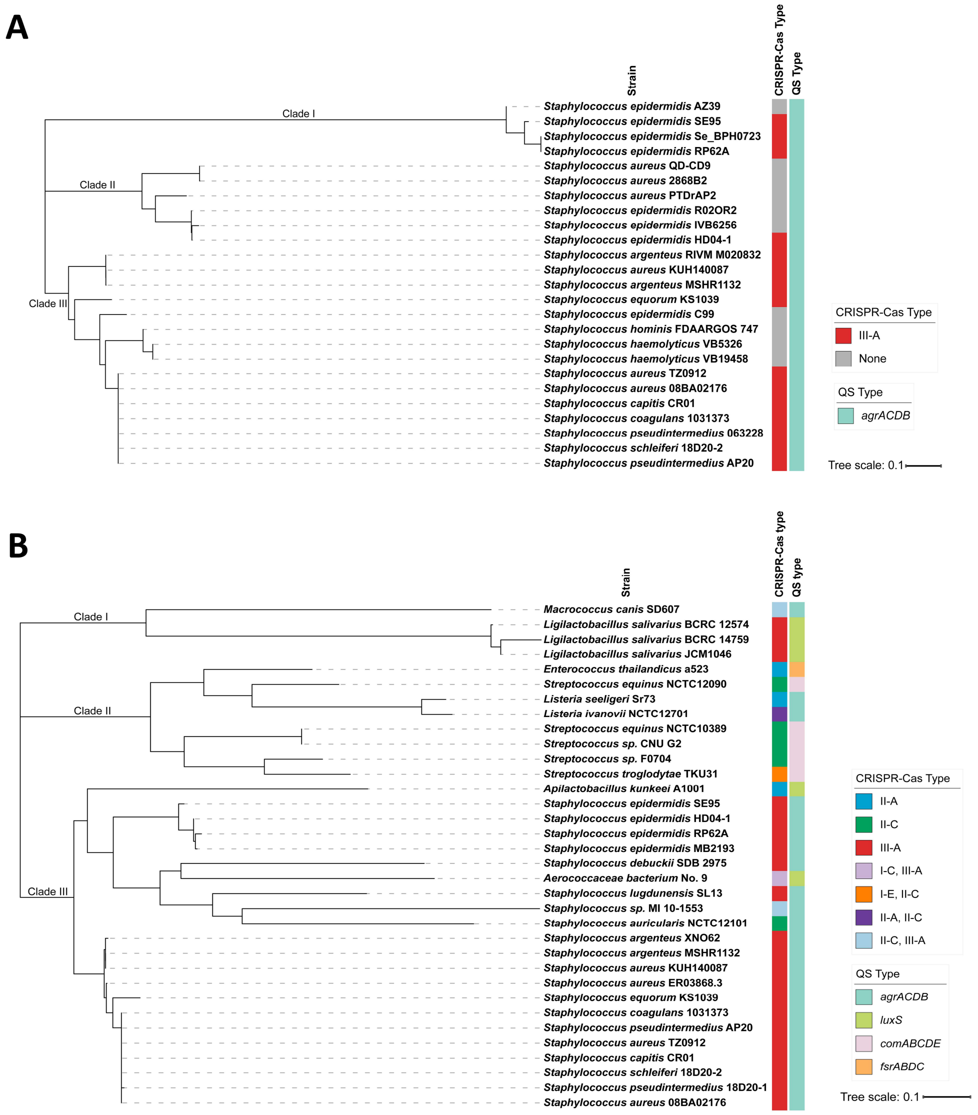


**Figure S6. Distribution of P*leader* and P*cas* sequences in Firmicutes. (A and B)** Phylogenetic trees were constructed based on P*leader* (A) and P*cas* (B) homologous sequences. More related P*leader* and P*cas* homologs were also identified in Firmicutes that are not present in trees, and are reported in Table S3 and S4, respectively.
